# Complete pan-plastome sequences enable high resolution phylogenetic classification of sugar beet and closely related crop wild relatives

**DOI:** 10.1101/2021.10.08.463637

**Authors:** Katharina Sielemann, Boas Pucker, Nicola Schmidt, Prisca Viehöver, Bernd Weisshaar, Tony Heitkam, Daniela Holtgräwe

## Abstract

**Background:** As the major source of sugar in moderate climates, sugar-producing beets (*Beta vulgaris* subsp. *vulgaris*) have a high economic value. However, the low genetic diversity within cultivated beets requires introduction of new traits, for example to increase their tolerance and resistance attributes – traits that often reside in the crop wild relatives. For this, genetic information of wild beet relatives and their phylogenetic placements to each other are crucial. To answer this need, we sequenced and assembled the complete plastome sequences from a broad species spectrum across the beet genera *Beta* and *Patellifolia*, both embedded in the Betoideae (order Caryophyllales). This pan-plastome dataset was then used to determine the wild beet phylogeny in high-resolution.

**Results:** We sequenced the plastomes of 18 closely related accessions representing 11 species of the Betoideae subfamily and provided high-quality plastome assemblies which represent an important resource for further studies of beet wild relatives and the diverse plant order Caryophyllales. Their assembly sizes range from 149,723 bp (*Beta vulgaris* subsp. *vulgaris*) to 152,816 bp (*Beta nana*), with most variability in the intergenic sequences. Combining plastome-derived phylogenies with read-based treatments based on mitochondrial information, we were able to suggest a unified and highly confident phylogenetic placement of the investigated Betoideae species.

Our results show that the genus *Beta* can be divided into the two clearly separated sections *Beta* and *Corollinae*. Our analysis confirms the affiliation of *B. nana* with the other *Corollinae* species, and we argue against a separate placement in the *Nanae* section. Within the *Patellifolia* genus, the two diploid species *Patellifolia procumbens* and *Patellifolia webbiana* are, regarding the plastome sequences, genetically more similar to each other than to the tetraploid *Patellifolia patellaris*. Nevertheless, all three *Patellifolia* species are clearly separated.

**Conclusion:** In conclusion, our wild beet plastome assemblies represent a new resource to understand the molecular base of the beet germplasm. Despite large differences on the phenotypic level, our pan-plastome dataset is highly conserved. For the first time in beets, our whole plastome sequences overcome the low sequence variation in individual genes and provide the molecular backbone for highly resolved beet phylogenomics. Hence, our plastome sequencing strategy can also guide genomic approaches to unravel other closely related taxa.

## Background

As the crop plant *Beta vulgaris* (sugar beet) has a high economic value (1), continuous crop development is essential to enhance stress tolerances and resistances against pathogens. The White Silesian Beet provided the germplasm for sugar beet (2) and is a derivative of wild sea beet (*B. vulgaris subsp. maritima*). This leaves, similar to the situation in many domesticated crops, only a narrow genetic base for sugar beet breeding (3). Additionally, early sugar beet breeding has focused mainly on increasing yield. This caused strong domestication bottle necks and removed many useful traits that may benefit plant fitness (3,4). The higher genetic variation in crop wild relatives of sugar beet offers potential that might be harnessed to introduce desired traits. Thus, giving insight into the genomic basis of wild beets is progressively moving into the focus of beet breeding research (5,6).

Phylogenetically, wild beets belong to the Betoideae (order Caryophyllales) and are separated in the genera *Beta* and *Patellifolia* (7,8), with all cultivated beets belonging to the genus *Beta* (9). The genus *Beta* is then further subdivided into at least two sections, *Beta* and *Corollinae*. In general, the section *Beta* is widespread across Western Europe, whereas *Corollinae* species are generally distributed across the eastern Mediterranean area and South-West Asia (1,8) (Additional file S1A). Despite the long history of different systematic treatments (1,7–12), the phylogenetic relationships of *Beta* and *Patellifolia* species are still a matter of ongoing debate (as reviewed in (7)). Especially the subdivision of the genus *Beta* into three sections (*Beta*, *Corollinae*, and *Nanae*) is discussed, with the pending suggestion to integrate *B. nana* into the section *Corollinae*, hence disbanding the section *Nanae* (7,8,13). Similarly, as the *Beta* section *Corollinae* harbors a highly variable polyploid/hybrid complex, including di-, tri-, tetra-, penta-, and hexaploid forms, the species boundaries are far from resolved. Regarding the sister genus, there is still an ongoing discussion on whether the morphologically variable *Patellifolia* comprise three distinct species (*P. patellaris*, *P. procumbens*, and *P. webbiana*) or only two or even one (8,10,12). Resolving the unclear wild beet relationships may inform beet improvement programs and contribute to the development of new, better equipped beets.

The plastome is well-suited for the reconstruction of phylogenies due to high structural conservation, a conserved evolutionary rate, uniparental inheritance, and high abundance of DNA across all species (14,15). Historically, systematic information was obtained from plastome sequence restriction site variants, inversions, single nucleotide variants (SNVs), or spacers in single genes. Although this has led to a range of wild beet phylogenies resolving relationships on the level of genera and sections, these are often based on only a few species and contain collapsed branches due to low genetic variation. In contrast, the investigation of whole plastome sequences may enhance the resolution of phylogenetic relationships (14,16–18). As gene sequences and intergenic regions can be included and combined, whole plastome sequence analyses enable the detection of well-supported phylogenetic relations on the species- and even on the accession level. Thus, plastid genomics may offer a route to clarify many of the pending questions regarding the wild beet phylogeny.

The plastome sequence of most angiosperms comprises a total of 79 protein-coding genes, 4 rRNA genes, and 30 tRNA genes (19). The quadripartite structure is characteristic for plastome sequences comprising a large (LSC) and a small single-copy region (SSC) as well as two inverted repeats (IRs) (20,21), all contributing to a total length of 120 kb to 210 kb (20). This difference in size can be mainly attributed to the IRs that range from 6 kb to 76 kb in length (21–23). The relative orientation of the SSC between the IRs differentiates two structural variants which occur simultaneously in a single cell and might have been previously mistakenly annotated as differences between species (24).

The Caryophyllales, including *B. vulgaris*, contain canonical plastomes, harboring all hallmarks typical for angiosperm plastome sequences as described above (25,26). For the wild beet species, until now, no plastome assembly is available, and of our investigated species, only the plastome sequence of *B. vulgaris* subsp. *vulgaris* was published previously (27). A detailed, plastome- and mitogenome-based evolutionary positioning of species outside of the section *Beta* is still missing but needed to answer some of the unresolved issues in beet systematics.

Here, we resolve the phylogenetic relationships within the Betoideae at high resolution through genome-wide comparison based on complete plastome assemblies and reads from both, the plastome and the mitogenome. Eleven different *Beta* and *Patellifolia* members, spanning the previously neglected plastome sequences of the *Corollinae* section and the *Patellifolia* genus, are included in our analyses. For this, whole plastome sequences of up to two accessions per species are sequenced, assembled, and compared. This novel contribution to the Betoideae pan-plastome intends to clarify the phylogenetic relationships of wild beets on a species-level and provides an important resource for further studies of beet wild relatives.

## Results

Our pan-plastome dataset comprises 18 different accessions, including a biological replicate of *B. corolliflora*, which leads to 19 plastome assemblies in total (see Methods, Table 2). To provide a basis for comparative plastome analysis, all plastome sequences were fully assembled. Out of those, 17 were split into three scaffolds (LSC, SSC, and IR), apart from Bmar1 (four scaffolds) and Bnan2 (six scaffolds). Collapsed IR regions were confidently identified in all plastome assemblies based on a doubled average read coverage in comparison to the single copy regions as well as a gene content that is characteristic and expected for the IR region. Average read coverage and assembly length are shown in Table 1. The distribution of these values is shown in Additional file S1B. Circular and linear plots of a representative selection of plastome assembly sequences are provided in Additional file S1C.

**Table 1:**
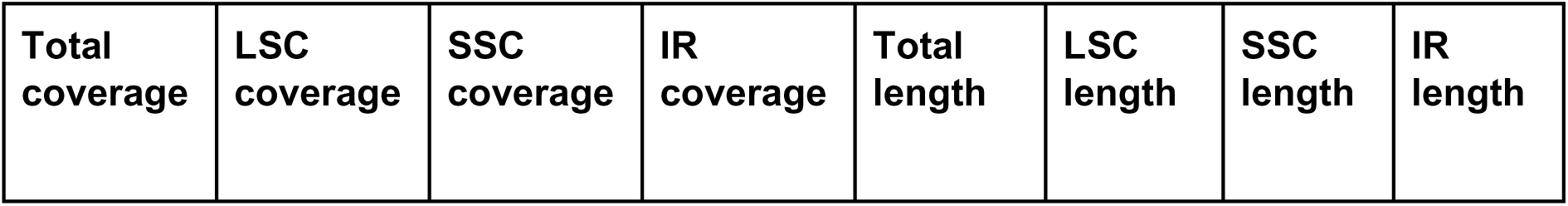
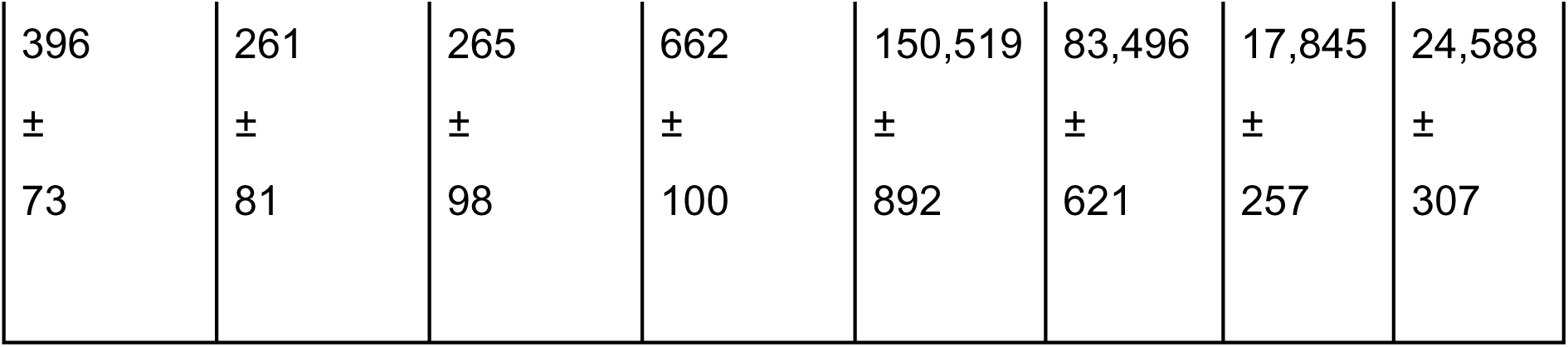
Plastome assembly statistics. . Average read coverage values and assembly lengths (in bp) for each region and the complete assemblies are shown. Abbreviations: LSC = Long single copy region; SSC = Short single copy region; IR = Inverted repeats.

| Total coverage | LSC coverage | SSC coverage | IR coverage | Total length | LSC length | SSC length | IR length |
| --- | --- | --- | --- | --- | --- | --- | --- |
| 396 | 261 | 265 | 662 | 150,519 | 83,496 | 17,845 | 24,588 |
| ± | ± | ± | ± | ± | ± | ± | ± |
| 73 | 81 | 98 | 100 | 892 | 621 | 257 | 307 |

**Table 2:** Abbreviation, species name, accession number, genus, and section of the investigated accessions of the Betoideae subfamily.

| Abbreviation | Species name | Accession number | Genus | Section |
| --- | --- | --- | --- | --- |
| Bmar1 | *Beta vulgaris* subsp. *maritima* | BETA 1101 | *Beta* | *Beta* |
| Bada | *Beta vulgaris* subsp. *adanensis* | BETA 1233 | *Beta* | *Beta* |
| Bmar2 | *Beta vulgaris* subsp. *maritima* | BETA 2322 | *Beta* | *Beta* |
| Bvul | *Beta vulgaris* subsp. *vulgaris* | KWS 2320 | *Beta* | *Beta* |
| Bptu | *Beta patula* | BETA 548 | *Beta* | *Beta* |
| Bmca | *Beta macrocarpa* | BETA 881 | *Beta* | *Beta* |
| Bnan1 | *Beta nana* | BETA 546 | *Beta* | *Corollinae*, formerly *Nanae* |
| Bnan2 | *Beta nana* | BETA 570 | *Beta* | *Corollinae*, formerly *Nanae* |
| Bint1 | *Beta intermedia* | BETA 431 | *Beta* | *Corollinae* |
| Bint2 | *Beta intermedia* | BETA 923 | *Beta* | *Corollinae* |
| Bcor1 | *Beta corolliflora* | BETA 408 | *Beta* | *Corollinae* |
| Bcor2 | *Beta corolliflora* | BETA 408 | *Beta* | *Corollinae* |
| Bmrh1 | *Beta macrorhiza* | BETA 830 | *Beta* | *Corollinae* |
| Bmrh2 | *Beta macrorhiza* | BETA 576 | *Beta* | *Corollinae* |
| Blom | *Beta lomatogona* | BETA 674 | *Beta* | *Corollinae* |
| Pweb | *Patellifolia webbiana* | BETA 526 | *Patellifolia* | - |
| Ppro | *Patellifolia procumbens* | BETA 951 | *Patellifolia* | - |
| Ppat1 | *Patellifolia patellaris* | BETA 534 | *Patellifolia* | - |
| Ppat2 | *Patellifolia patellaris* | BETA 892 | *Patellifolia* | - |

Comparing the plastome assemblies of all *Beta* vs. *Patellifolia* species, the average length of the four *Patellifolia* plastome sequences (avg. 151,621 bp) is higher than for the 15 *Beta* plastome sequences (avg. 150,225 bp). This length difference can be mainly assigned to the LSC (avg. *Beta* 83,401 bp; avg. *Patellifolia* 83,853 bp) and to the IRs (avg. *Beta* 24,435 bp; avg. *Patellifolia* 25,166 bp). However, the SSC is longer in *Beta* plastome sequences (avg. 17,954 bp) when compared to the plastome sequences of all *Patellifolia* accessions (avg. 17,437 bp).

Interestingly, plastome assemblies of *B.* section *Corollinae* (avg. 36.67 %) show a higher GC content when compared to *B.* section *Beta* plastome sequences (avg. 35.81 %). The total length of *B.* section *Corollinae* plastome sequences (avg. 150,504 bp from nine species) is higher in comparison to *B.* section *Beta* plastome assemblies (avg. 149,808 bp from six species). This length difference is visible for all regions of the plastome sequence (LSC, SSC and IRs).

The final plastome assemblies were subsequently annotated and the alignment identity of all regions included in the phylogenetic analysis was assessed for gene regions and intergenic regions, respectively (Figure 1). The alignment identity is significantly higher for gene sequences when compared to intergenic regions. This significant difference was obtained when amaranth, quinoa, and spinach were included as outgroups (Additional file S2A) (avg. gene/intergenic regions 90.73/83.55 %; Mann-Whitney-U test; p 2e-10) as well as without outgroup reference sequences (Additional file S2B) (avg. gene/intergenic regions 97.26/94.93 %; Mann-Whitney-U test; p ≈ 1e-09). *Rrn* genes show high similarity among all plastome genes, whereas *ycf1* and *rpl22* show the greatest variance between all investigated accessions. The intergenic region between the genes *ycf4* and *cema* contributes most to the differences in the alignment.

**Figure 1:**
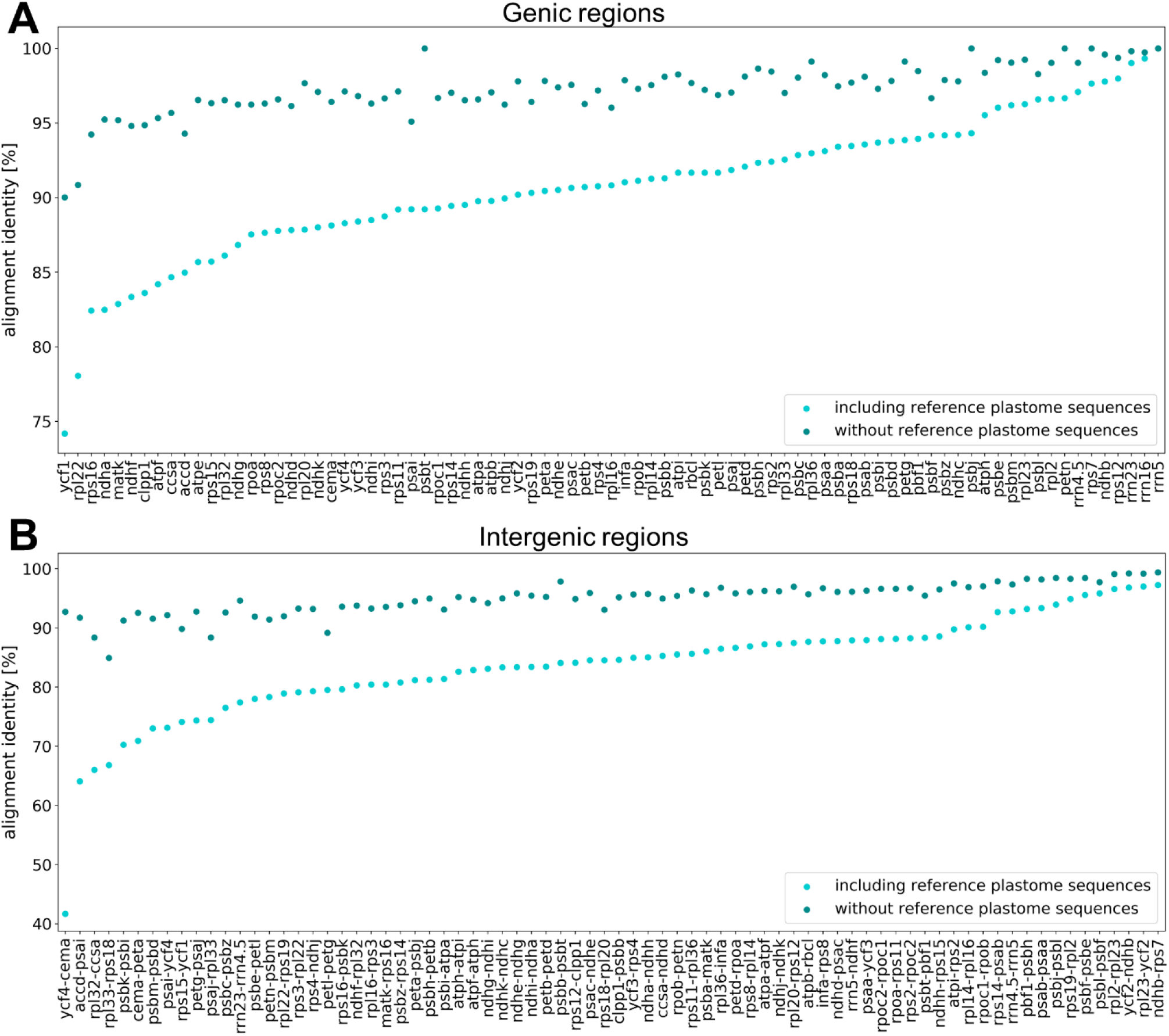
Alignment identities of gene sequences (A) and intergenic regions (B). The sequences (x-axis) are ordered based on the alignment identity represented by the y-axis. The alignment identity including amaranth, quinoa, and spinach as outgroup (light blue) as well as the alignment identity without the reference plastome sequences (dark blue) are shown.

The distributions of SNVs (Figure 2A) and InDels (Figure 2B) throughout the plastome sequences were further investigated. InDels are mostly absent from gene regions and SNVs are in general more frequent than InDels. Further, some clear hotspots of SNVs and InDels can be detected in the intergenic regions *psbK-psbI*, *ycf4-cema*, *rpl33-rps18*, *psaj-rpl33* and *rpl32-ccsa*.

**Figure 2:**
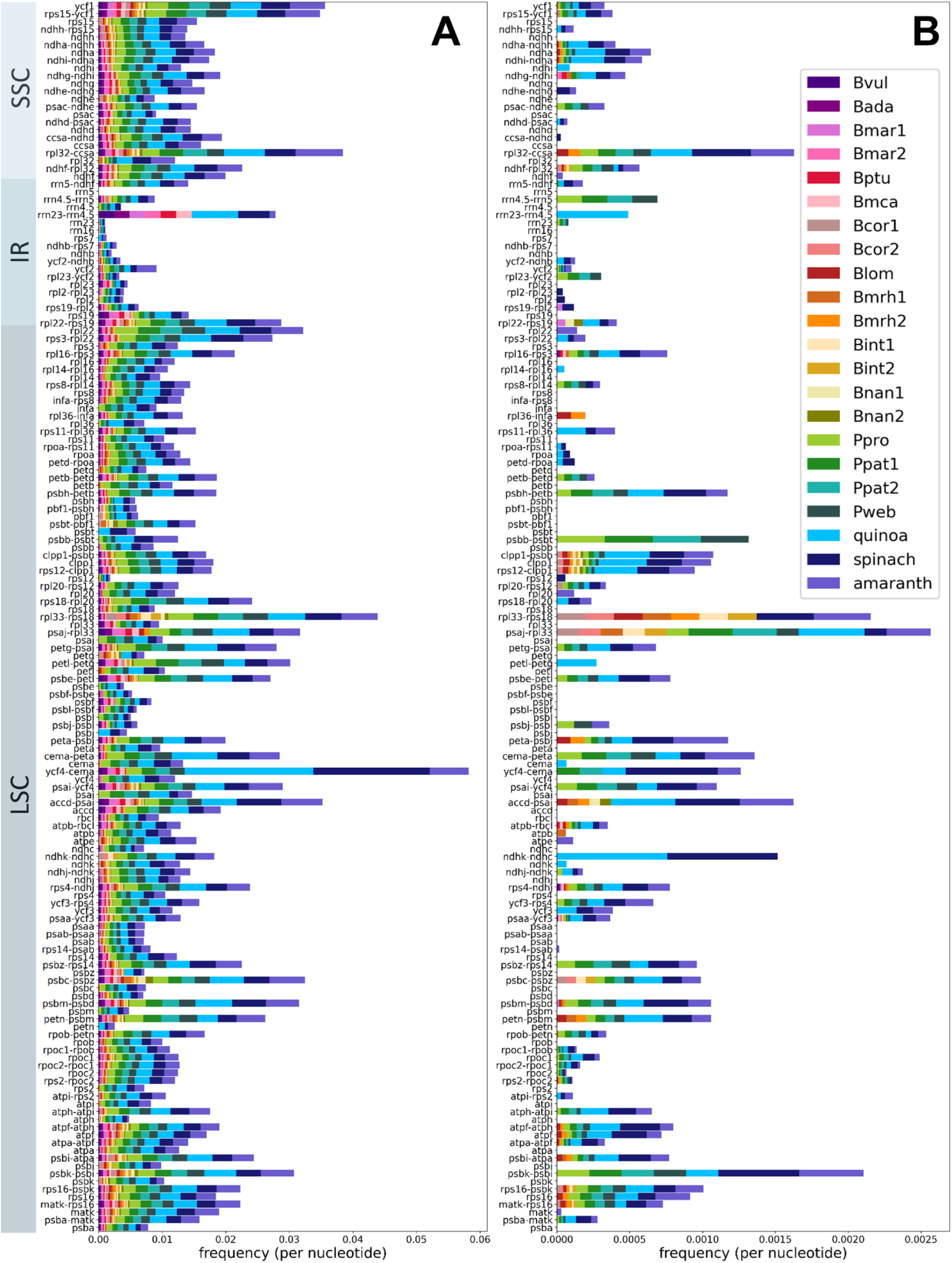
Number of SNVs and InDels in the assembled plastome sequences. Number of SNVs (A) and number of InDels (B) in the sequence alignments normalised by the length of the respective gene sequence/intergenic region and by the number of species/accessions. The gene names and intergenic regions on the x-axis are ordered based on the arrangement in the plastome assembly. Amaranth, quinoa, and spinach were included as outgroup.

As InDels with a length, which is a multiple of three, do not influence the reading frame (28), we expected that the proportion of these InDels (which are multiples of three) is higher in gene regions compared to the proportion in intergenic regions. Indeed, we observed that 43.4 % of the InDels in gene regions were a multiple of three, whereas this applies to only 29.6 % of the InDels in intergenic regions (Fisher’s exact test; p ≈ 3e-8; Figure 3 [arrows]).

**Figure 3:**
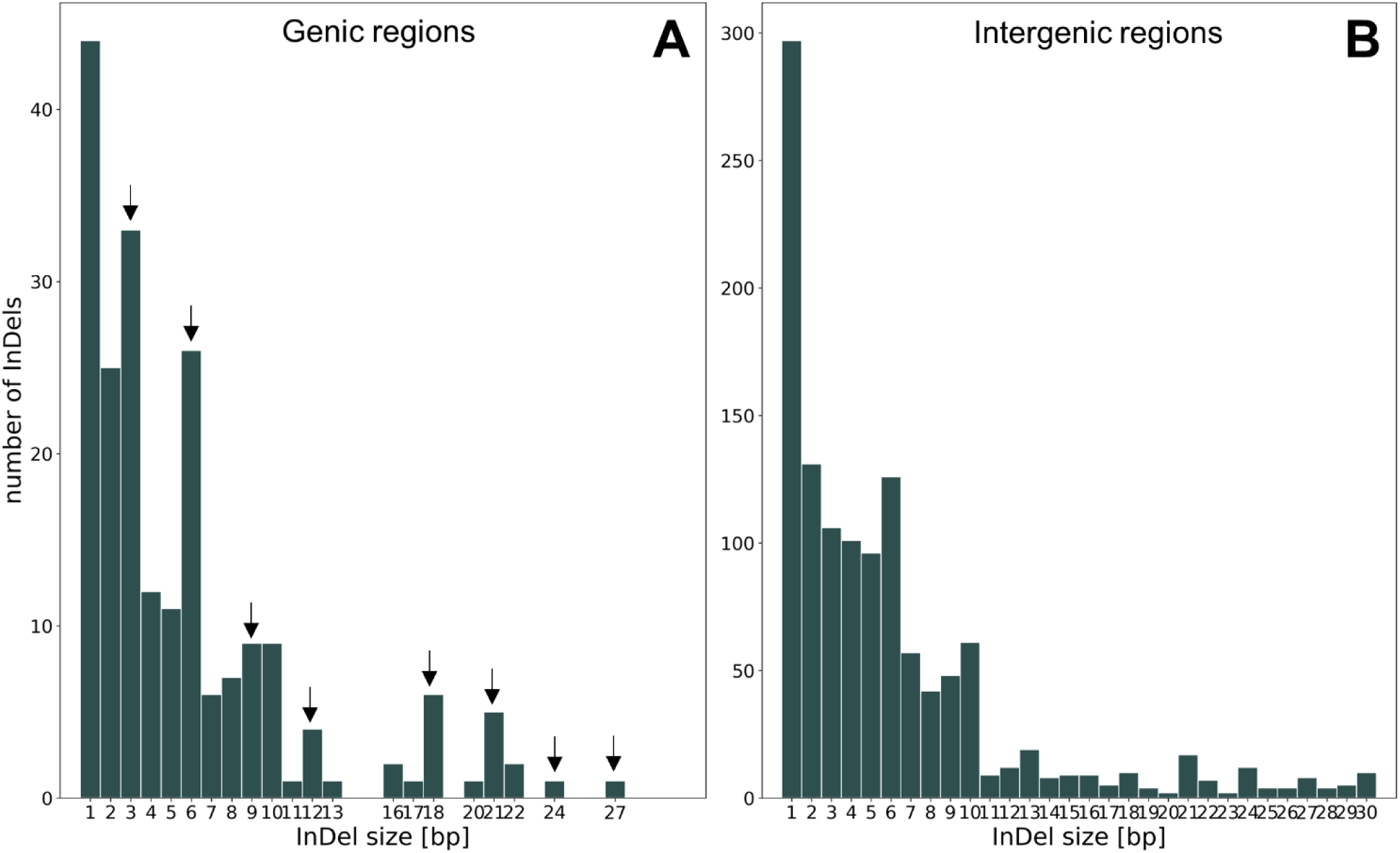
Number of InDels based on size in gene sequences (A) and intergenic regions (B). The arrows represent InDels with a size of a multiple of three. Please mind the variable y-axis.

Phylogenetic relations of the Betoideae subfamily were inferred from colored de Bruijn graph-based splits (Figure 4) as well as by an alignment-based maximum likelihood (ML) analysis (Figure 5). Mitochondrial and plastid read derived kmers were used to calculate phylogenetic splits, and annotated gene sequences as well as intergenic regions derived from the plastome assemblies were used for the ML analysis. A clear separation of *Patellifolia*, *B.* section *Beta* and *B.* section *Corollinae* samples is visible in all four phylogenetic trees. In comparison to the ML tree based on fully assembled plastome sequences (Figure 4C, black), the tree based on splits derived from the same dataset (fully assembled plastome sequences) (Figure 4C, dark green) shows only one difference among the *B. intermedia* and *B. corolliflora* accessions. The phylogenetic relationships derived from kmers show a few additional differences (Figure 4C, light green and pink). These differences comprise for example the assignment of the four *Patellifolia* accessions/species to a clade consisting of both *P. patellaris* accessions and a separate clade formed by *P. procumbens* and *P. webbiana* for the assembly-based phylogenies (Figure 4C, black and dark green), whereas the kmer based phylogenies show a separate clade for *P. webbiana* and a second clade comprising the other three *Patellifolia* accessions/species (Figure 4C, light green and pink). The calculation of the weighted F1 score, the weighted symmetric set distance and the Robinsons-Foulds distance shows that there is a high identity between all splitstree results (based on cp_reads, mt_reads and cp_assemblies) (Additional file S1D). The usage of different input data formats (reads vs. assemblies) has a larger impact than the usage of different datasets (chloroplasts vs. mitochondria) meaning that these splitstree results show a higher divergence.

**Figure 4:**
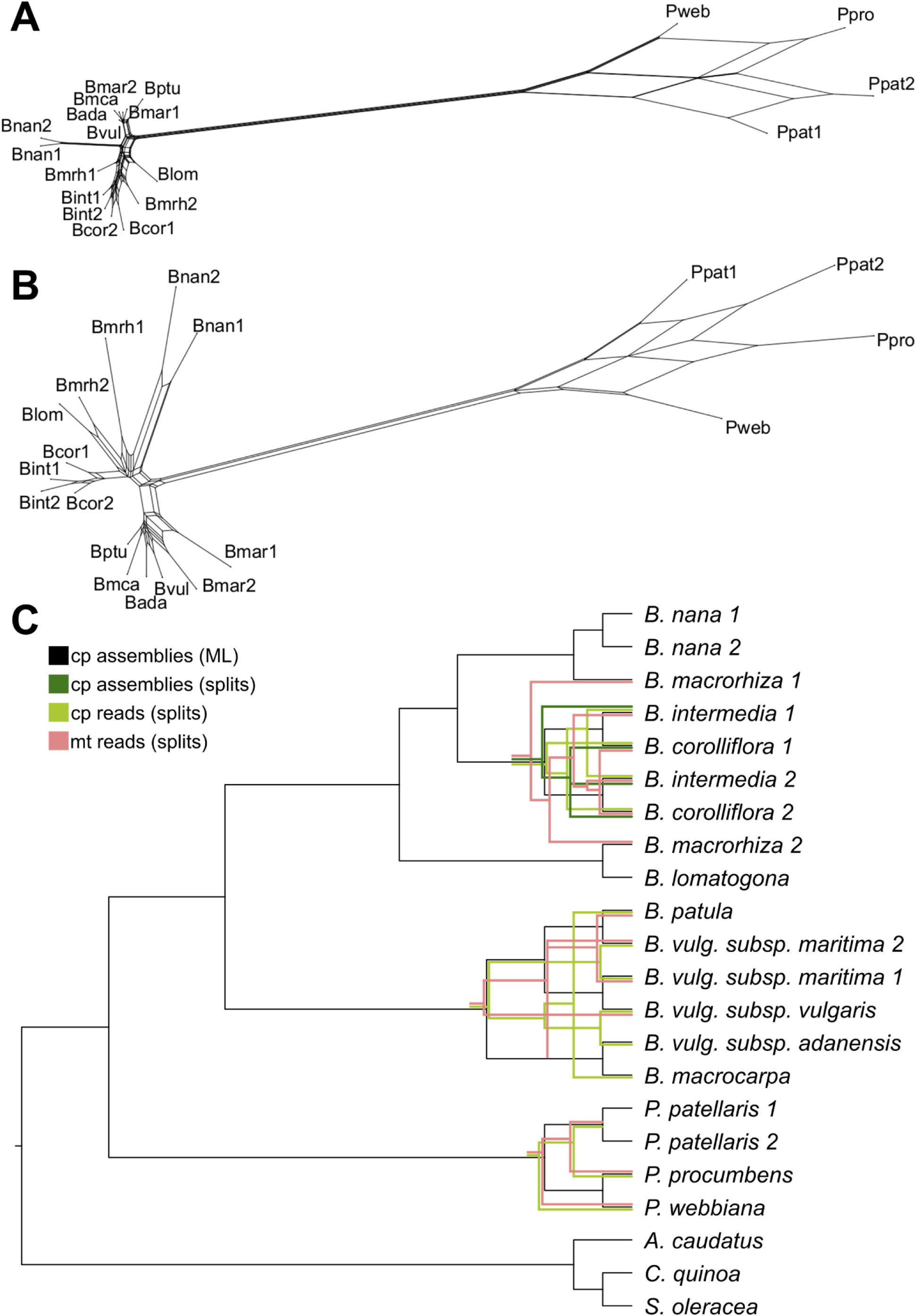
Phylogenetic relationships of 18 different *Beta*/*Patellifolia* accessions derived from different strategies and datasets. Kmer-based trees were constructed using raw sequencing reads (A: mitochondrial reads, B: chloroplastic reads) as well as the final chloroplast assemblies as input (not shown as only used for comparison). The splitstree results (green and pink) were then compared to the ML analysis (black) (C). Discordance between the phylogenetic trees is shown in the respective color. (Abbreviations: cp=chloroplast; mt=mitochondria).

**Figure 5:**
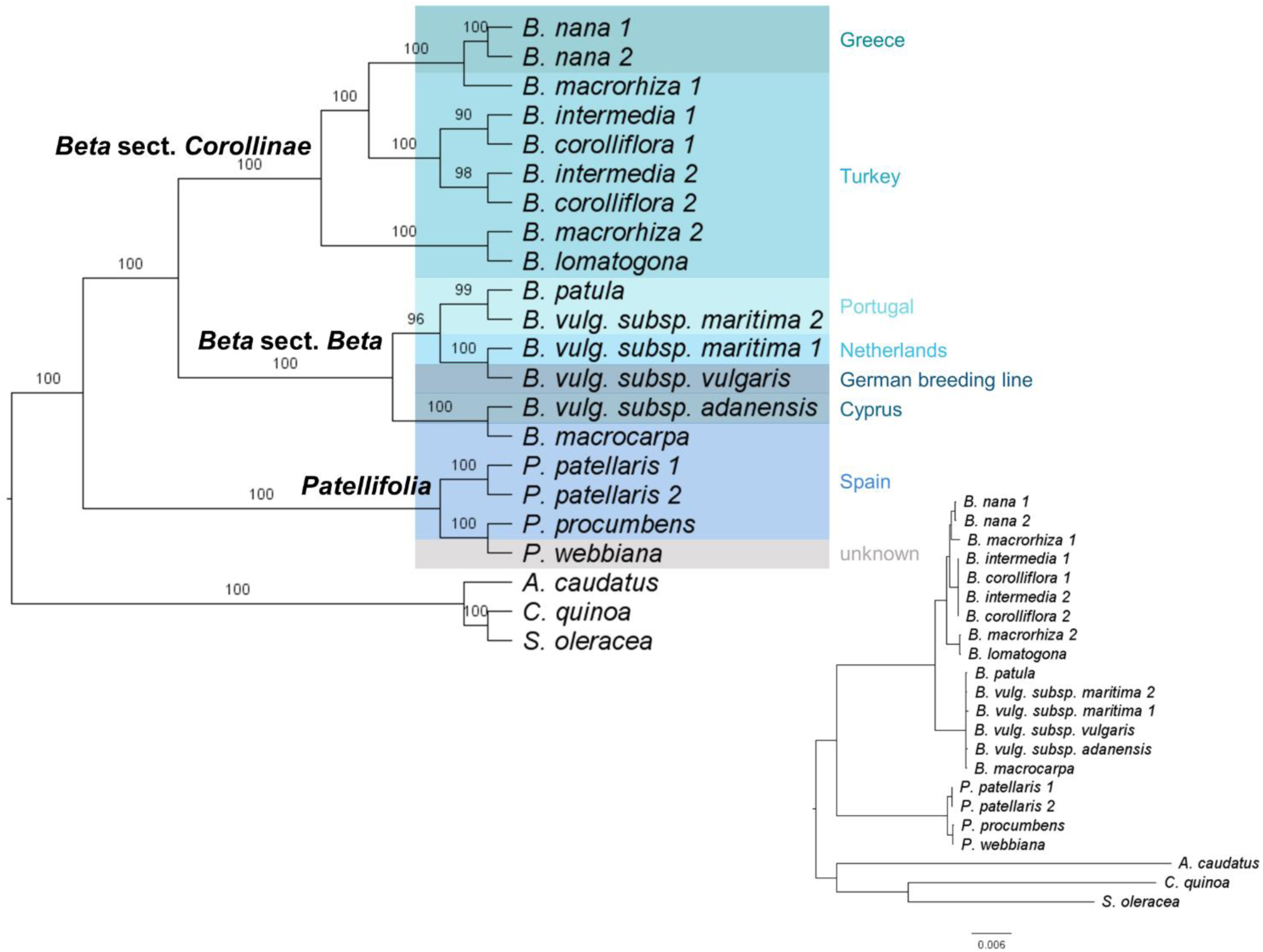
Plastome phylogeny for Betoideae. The tree can be divided into three groups: *B.* section *Corollinae* (8 accessions), *B.* section *Beta* (6 accessions), and *Patellifolia* (3 accessions). The plastome sequences of three Caryophyllales species (amaranth, quinoa, and spinach) were used as outgroup. Bootstrap support values are shown above each branch. The resulting phylogeny is based on the variation in 83 genes and 76 intergenic regions from the plastid genomes of 18 accessions and species (plus outgroup). Different background colors represent the sampling location and the origin of the breeding line (Bvul), respectively. Inset: Actual branch lengths based on the ML analysis.

For the ML-based tree (Alignment sites / patterns: 216442 / 2441; Gaps: 0.44 %; Invariant sites: 86.73 %), the phylogenetic relationships on the species level are highly supported (high bootstrap values) (Figure 5). A phylogenetic tree based on the diagnostic set of 53 gene sequences and intergenic regions matches this phylogeny (Additional file S1E).

To investigate the contribution of different regions to the phylogeny, in addition to the whole plastome sequences (genic and intergenic regions combined), sequence matrices for (I) all gene regions, (II) all intergenic regions, (III) whole coding sequences, (IV) first and second codon position and (V) third codon position only were constructed and used for the inference of phylogenetic relationships. Even though the topology of the phylogeny derived from whole coding sequences is highly similar (Additional file S1F), especially for the codon position-based matrices, alternative branches can be observed. However, substantially more nodes are only poorly supported with bootstrap values up to below 20.

The alignment of the 18s rRNA gene sequence for *B. corolliflora* (Bcor1) as representative for the *Corollinae* and *B. vulgaris subs. vulgaris* as member of the section *Beta* revealed not a single genomic difference and therefore no signal which could be used to resolve their phylogenetic relations.

The results presented here show that especially for closely related species of the same subfamily, a higher number of gene sequences and more variable intergenic regions provides greater phylogenetic resolution.

## Discussion

### Our proposed beet whole-plastome phylogeny is superior to single gene phylogenies

Here, we present the plastome sequence assemblies of 18 accessions covering most of the species’ diversity within the beet genera *Beta* and *Patellifolia* and representing an important resource for future studies. All newly generated plastome sequences are highly similar including 79 protein coding genes and four rRNA genes distributed across a mean length of 150,519 bp (± 892bp). A previously published *B. vulgaris* subsp. *vulgaris* (KR230391) plastome sequence comprises a length of 149,722 bp (27). This is almost identical to our Bvul assembly which differs by only 1 bp in length (149,723 bp). This difference occurs in a stretch of five or six guanines in the IR region between the genes *rrn23* and *rrn16* – either a sequencing error or a biological difference. As expected, all Betoideae assemblies show high similarity as all angiosperm plastomes are highly conserved and the species investigated here are closely related, while most differences are located in intergenic regions.

Between the cultivated beet and wild beet accessions, most chloroplast genes are highly conserved, one example being *rpl2* with only a low number of polymorphisms (Figure 1) (22). The intergenic regions are significantly less conserved containing more InDels and are therefore more suitable for phylogenies on a lower taxonomic level (22). Among the beet plastomes, *ycf1* is the most informative genic region (Figure 1A), which was also detected in other plant groups, such as the Tropaeolaceae, the orchids, and the Malvales (29–31). Additionally, the *rpl22*, *matk*, *clpP1*, *ycf2*, *psaC* and *ndhF* genes were reported to be highly divergent (29,31,32), which is mainly consistent with our findings. Here, the rrn-genes, *ndhB* (already classified as gene with low divergence (29,31)), *rps12* and *rps7* can be found among the gene loci with lower variance among the investigated species.

Plastomes in general show very similar sequences with most differences occurring in non-coding regions (33). For beet and wild beets, the most informative intergenic regions are *ycf4-cema*, *accd-psai* and *rpl32-ccsa* (Figure 1B). However, for *ycf4-cema* the high difference between ‘reference’ and ‘without reference plastome-’ alignment identity should be noticed. For the Malvales, the regions *psaB-psaA*, *psbF-psbE*, *rpl2-rpl23* and *ndhH-ndhA* were identified as lowly divergent regions (31). We confirmed these for beets, except for the latter (*ndhH-ndhA*) that showed higher divergence among the wild beet plastomes. In contrast, the intergenic regions with the highest divergence in the Malvales (*ndhD-ccsA* and *rps19-rpl2*) (31) accounted for less differences among the wild beets. Nevertheless, the percent identity values among all intergenic regions are relatively similar in beets, especially when excluding the polymorphisms in the outgroup reference genomes.

As high variability among the investigated sequences is required to resolve phylogenetic relations of closely related species (34), the retrieved plastome sequences from beets and wild beets provide an excellent resource to approach systematic treatment of the Betoideae. To further increase sequence variability, we also integrated the more diverged intergenic regions into the analysis.

In addition to the plastome-inferred ML-based phylogeny, a mitogenome- and plastome-inferred read-based phylogenetic tree was constructed based on phylogenetic splits. The resulting trees can be considered reliable due to three distinct, robust properties: (I) all splits networks show a tree-like appearance (Figure 4A, 4B), (II) the usage of different kmers leads to the same tree and (III) the usage of the geometric mean leads to the removal of samples in case no kmer occurs in the sample and both, geom and geom2 lead to the same results (with the exception of k=11, which results in a cloud-shaped network).

Differences between the phylogenetic trees derived from different strategies and datasets (Figure 4C) may be explained by chance and/or by the use of the method (all splitstree results show high identities as indicated by all comparison metrics (Additional file S1D) while differences are larger when using different strategies [reads vs assemblies] in comparison to different datasets [chloroplasts vs. mitochondria]) or by real differences in the biological nature of mitochondria and chloroplasts. Mitochondrial DNA shows a low nucleotide substitution rate when compared to chloroplast DNA (25,35). Reasons for this may be recent species hybridization or incomplete lineage sorting (36,37). Therefore, the mitogenome seems to be mostly useful at higher taxonomic levels (25) and might not be the most suitable system for the beet and wild beet accessions investigated in this study. In summary, the trees based on plastome assemblies (ML and splits) are likely the most reliable as the same phylogenetic relations are the outcome of different established strategies including the widely used ML method.

Compared to our pan-plastome assembly, the information derived from individual genic and intergenic regions are insufficient to fully resolve the beet phylogeny, highlighting the power of our plastome approach:

1. Investigation of specific regions of the plastome sequence (genic and intergenic regions, coding sequences, codon positions) revealed a few alternative branches for the codon position-based sequence matrix (Additional file S1F). These are marked by short internal branch lengths due to the close relationships of the species within this subfamily. Nevertheless, these conflicting relationships are only weakly supported. This is explained by the lower genetic diversity and therefore insufficient phylogenetic signal when using a smaller amount of sequences and total sequence length.
2. An approach based on high-quality single-copy nuclear genes would require a minimum coverage of about 10x (25). Moreover, nuclear genes are often part of gene families and influenced by whole genome duplication events (14). Using our available data, nrDNA sequences were selected for the phylogenetic reconstruction. Especially the ITS and ETS regions were previously used for the investigation of phylogenetic relationships, but entire nrDNA repeats (18S-ITS1-5.8S-ITS2-26/28S) were also already assembled for multiple phylogenetic studies (15,38). Unfortunately, the assembled nrDNA sequences constructed here are not useful to infer confident phylogenetic relationships as the coverage is very low (1.9x - 8.5x) and intragenomic polymorphisms of different nrDNA repeat sequences might limit the reliability of the phylogeny (15). The low bootstrap values and the low coverage make this phylogenetic tree unreliable. Therefore, we do not show these results here. Another possible explanation, apart from the low coverages, is that the biparental nature of the nuclear genome may be problematic for the inference of phylogenetic relationships (14). Previous studies already suggest that phylogenies based on nrDNA and few selected plastid sequences only weakly support relationships (30). The 18s rRNA gene sequences of representatives of the sections *Corollinae* and *Beta* are completely identical containing no phylogenetic signal to separate them.

Summarizing, our plastome-derived phylogeny benefits from the incorporation of genic and intergenic regions as well as the ‘nature’ of the plastome itself (as described in the Background section). Despite the low available read coverage and the low genetic diversity within our beet dataset, this leads to a highly confident phylogenetic tree. Further, the use of higher alignment lengths and the use of nucleotides instead of amino acids are favored to construct well supported phylogenies (39). Therefore, we conclude that for resolving the relationships of cultivated and wild beets, our whole-plastome-based approach is the most reliable.

### Implications for the systematic placements within the Betoideae subfamily

With efforts tracing back half a century, resolving the phylogeny of the subfamily Betoideae has been already a major undertaking (1,8–11,40). However, in most cases only few selected sequence regions were targeted, leading to unresolved relations at shallower taxonomic levels or with focus on specific species or sections, i.e.:

1. In a study by Hohmann et al. (2006), the *Beta* species *B. vulgaris*, *B. corolliflora*, *B. nana* and *B. trigyna* were investigated using ITS, *trnL-trnF* spacer and *ndhF* sequences (10). Kadereit et al. (2006) provide a comprehensive analysis of a high number of different Betoideae species, finding that, *B.* section *Beta* was clearly separated from *B.* section *Corollinae*, which contained *B. nana*, *B. trigyna*, *B. macrorhiza*, *B. corolliflora* and *B. lomatogona*. As the analysis was based on ITS sequences comprising only 251 characters of which 147 were invariable, relations between species in both sections could not be resolved (8). Our study confirms the deep separation of the sections *Beta* and *Corollinae* and refines the resolution on the species level.
2. A recent comprehensive study by Romeiras et al. (2016) of phylogenetic relationships in the Betoideae is based on ITS and *matK*, *trnH-psbA*, *trnL* intron and *rbcL* sequences and investigates a high number of different species leading to the following main result: *Beta* and *Patellifolia* species are two clearly separated monophyletic groups (1). In total, three monophyletic lineages were identified: *B.* section *Beta* (*B. vulgaris* subsp. *vulgaris*, *B. vulgaris* subsp. *maritima*, *B. macrocarpa*, *B. patula*), *B.* section *Corollinae* (*B. nana*, *B. corolliflora*, *B. trigyna*) and *Patellifolia* (*P. patellifolia*, *P. procumbens* and *P. webbiana*). Exact relations within the Betoideae on a lower taxonomic level remain unclear as the branches are not well supported and collapsed. We also identify the three monophyletic groups as proposed and manage to resolve many of the previously collapsed branches.
3. Recently, Touzet et al. (2018) investigated the relationship of a wide range of *B. vulgaris* subsp. *maritima*, *B. macrocarpa* and *B. vulgaris* subsp. *adanensis* accessions based on a 3,742 bp alignment of plastome sequences and a 1,715 bp alignment of selected nuclear sequences (11). They find, based on a representative geographical sampling, that *B. macrocarpa* is a distinct lineage from the two investigated *B. vulgaris* subspecies. Despite this interesting finding, the suggested phylogeny did not focus on other important species and accessions of the Betoideae subfamily and might be further improved by the analysis of sequences with higher diversity to reach higher bootstrap values, which we achieved using intergenic and genic regions of the whole plastome sequences.

With our pan-plastome-informed datasets, we have been able to confirm many of the observations before and added an unprecedented resolution at the species-level. More in detail, we conclude that:

1. Among the section *Beta*, the plastome sequences of *B. patula*, *B. vulgaris* subsp. *vulgaris* and *B. vulgaris* subsp. *maritima* are highly similar as indicated by the slightly lower bootstrap values (96/100) for these three beets. As this section harbors wild beets in relatively close geographical proximity across the coastal Mediterranean area, the detected similarity can be explained by (natural) crossing and gene flow due to close geographical proximity or accidental cross-pollination during cultivation as wild beet and cultivated beet groups are easily cross-compatible (3). Further, *B. vulgaris* subsp. *vulgaris* and some *B. vulgaris* subsp. *maritima* accessions are even phenotypically highly similar (41). The phylogenetic relationships among species can also be influenced by the geographical distribution, mating systems and polyploidization (11). Allogamy and self-incompatibility are characteristics of *B. vulgaris* subsp. *maritima*, whereas *B. macrocarpa* and *B. vulgaris* subsp. *adanensis* are self-compatible leading to lower divergence and higher homozygosity. Cross-compatibility can lead to hybridization by facilitating gene flow between individual species (40), especially between *B. patula* and *B. vulgaris* subsp. *maritima*, which may explain the lower bootstrap support and the more unclear relations in the phylogenetic tree presented here. Although previous studies found low divergence between *B. vulgaris* subspecies (11), *B. vulgaris* subsp. *adanensis* seems to be clearly separated from the other subspecies in our analysis, possibly explained by the geographical distance to the other investigated samples.
2. We confidently assigned specific species, including *B. nana*, to the section *Corollinae*: *B. corolliflora, B. intermedia, B. macrorhiza, B. lomatogona a*nd *B. nana c*luster together and form this *s*ection (also suggested by (8)). Here, we particularly focused on the *B. s*ection *Corollinae* by analysing the plastome sequences of eight accessions from five different species plus a biological replicate of the eponymous species *B. corolliflora. B. nana*, which is endemic to Greece (1,8,42), was previously considered a separate *B.* section *Nanae*. Our results, however, combined with multiple other studies, clearly show that *B. nana* falls within the *B.* section *Corollinae*, which is distinct from *B.* section *Beta* (8,10,13). In addition to our plastome-based phylogenetic analysis, further genomic evidence points to high genomic similarities between *B. nana* and other *Corollinae*: For example, these species are marked by similar repeat accumulation profiles as shown for many individual transposable element types (43–46). Regarding plant characteristics, frost tolerance and seed hardiness are useful traits in the section *Corollinae*, including *B. nana*, but do not occur in any species of the section *Beta* (47). Thus, frost tolerance is specific to the *Corollinae* when compared to the *Beta* and *Patellifolia* species. These points lead to the classification of *B. nana* as a member of the *Corollinae*. Considering the highly variable polyploid/hybrid status complexes within the *Corollinae*, our plant set encompassed three diploids (*B. macrorhiza*, *B. lomatogona*, and *B. nana*), a tetraploid (*B. corolliflora*), as well as a pentaploid (*B. intermedia*). Although the hybrid status and parental contributions of the polyploids remain unresolved (1,40), we present convincing evidence that *B. intermedia* and *B. corolliflora* are closely related. Thus, our plastome sequence analysis brings new evidence supporting the hypothesis that *B. corolliflora* and *B. intermedia* belong to a highly variable polyploid hybrid complex (summarised by (7); Figure 5). The investigation of the whole genome sequences of these polyploid species may help to resolve these parental contributions.
3. Among the *Patellifolia* members, *P. procumbens* and *P. webbiana* can be phylogenetically distinguished: *Patellifolia* was previously classified as *B.* section *Procumbentes* and there is still an ongoing taxonomic debate whether *P. patellaris*, *P. procumbens* and *P. webbiana* can be considered as separate species. However, due to molecular and morphological traits, *Patellifolia* are now mostly considered a separate genus which is divided into three distinct species (8,10,12). The relationships among the *Patellifolia* species could not be resolved in previous studies (1). In the phylogenetic tree presented here, *P. procumbens* and *P. webbiana* seem to be closely related (however still distinguished with high support) and clearly separated from the two *P. patellaris* accessions. The branch lengths distinguishing *B. patula* and the *B. vulgaris* subsp. *vulgaris***/***maritima*, which are both considered separate species, are highly similar (0.0001). The same branch length separates *P. procumbens* and *P. webbiana*. Therefore, our phylogenetic analysis indicates that the three *Patellifolia* species are distinguishable on a molecular level.

Comparing the results presented here with earlier studies, the previous investigation of Betoideae was substantially extended and refined. The phylogenetic relationships were resolved in more detail and not only based on the monophyletic groups. This is especially important for the species of the *B.* section *Corollinae* which were investigated in depth. Using the whole plastome sequences, including intergenic regions, it was possible to further resolve the phylogenetic relationships with higher bootstrap support due to the extraction of higher sequence variance and phylogenetic signal within the subfamily.

## Conclusions

We provide 19 plastome assemblies for 18 different beet and wild beet accessions, which can also be re-used for future investigations of beets and other Caryophyllales species, and harnessed these to revisit systematic issues within the genera *Beta* and *Patellifolia*. This analysis advanced our understanding of the phylogenetic relationships of the subfamily Betoideae in four ways: I) Analysing sequences of intergenic regions of the whole plastome assemblies made it possible to reveal the phylogeny of closely related species with high reliability. Our phylogenetic tree shows a clear separation of the wild beet genera *Beta* and *Patellifolia*, as well as of the two sections *Beta* and *Corollinae*. II) *B. vulgaris* subsp. *adanensis* and *B. macrocarpa* can be clearly distinguished from *B. vulgaris* subsp. *vulgaris*, *B. vulgaris* subsp. *maritima* and *B. patula*. A clear split of *B. patula* from the two *B. vulgaris* subsp. (*B. vulgaris* subsp. *vulgaris* and *B. vulgaris* subsp. *maritima*) was not observed, likely due to the high sequence identity possibly explained by the close geographical proximity and the fact that these species are easily cross-compatible. III) All three *Patellifolia* species are clearly separated in our phylogenetic analysis, while *P. procumbens* and *P. webbiana* are more closely related to each other than to *P. patellaris*. These results, including the investigation of the branch lengths, point to a molecular separation within the *Patellifolia* species. IV) Finally, the taxonomic classification of *B. nana* as a member of the *Corollinae* was further supported.

## Methods

### Plant material, genomic DNA extraction, and DNA sequencing

Seeds of Betoideae species were obtained from KWS Saat SE, Einbeck, Germany (*B. vulgaris* subsp. *vulgaris* genotype KWS 2320) and from the Leibniz Institute of Plant Genetics and Crop Plant Research Gatersleben (IPK), Germany (all other accessions with accession numbers listed in Table 2 and Additional file 2). The material of the KWS SaaSE, Einbeck and IPK Gatersleben was transferred under the regulations of the standard material transfer agreement (SMTA) of the International Treaty.

Apart from *B. vulgaris* subsp. *vulgaris*, 17 other *Beta* and *Patellifolia* accessions shown in Figure 6 were analysed. The exact sampling location of the investigated accessions was extracted from the GBIS/I (Genebank information system; IPK) (48) (Figure 6).

**Figure 6:**
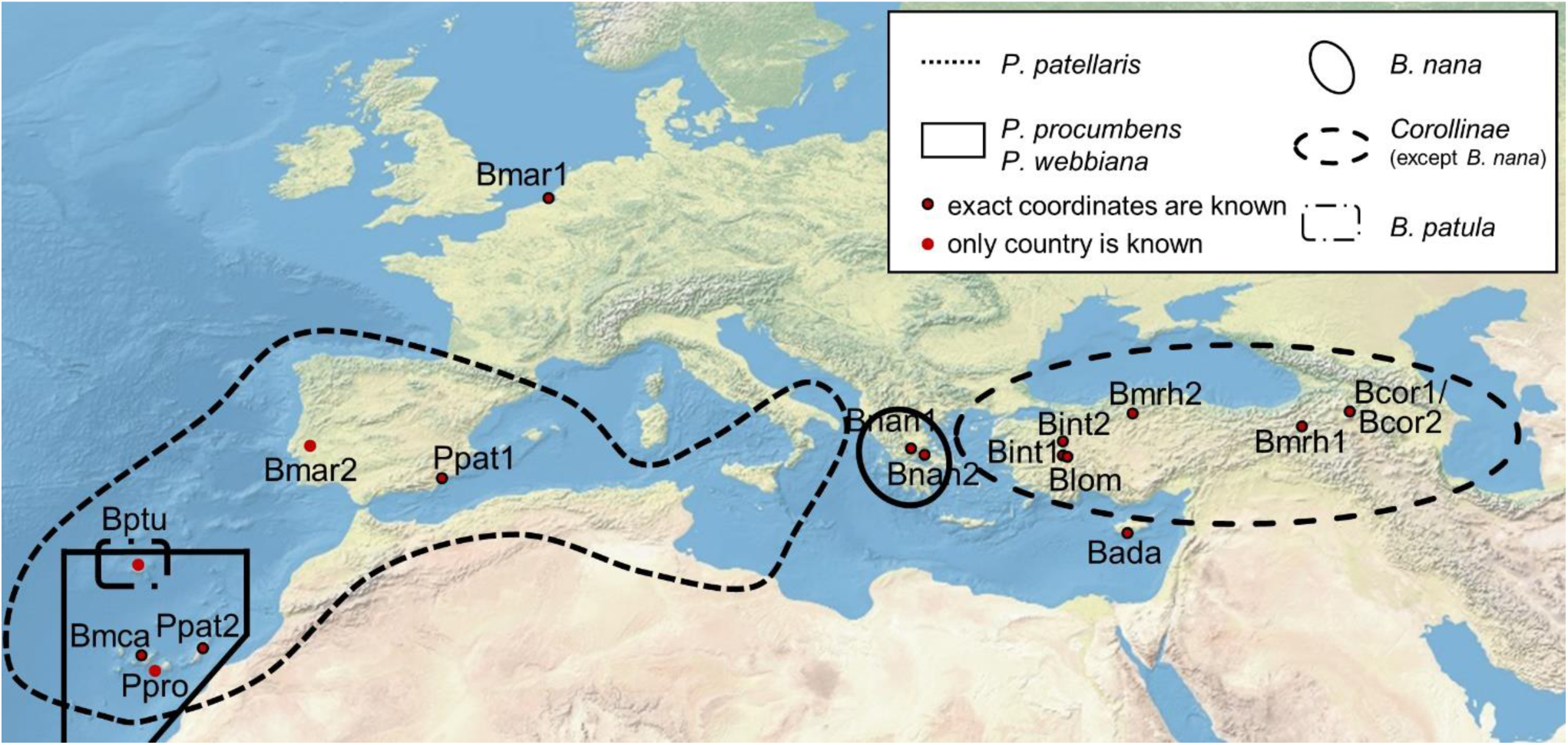
Geographic distribution of the *Beta* and *Patellifolia* species. The exact sampling locations of the investigated species are shown. The black lines represent the distribution area of the respective species and sections (see legend). The distribution areas of *B. vulgaris* subsp. *maritima*, *B. vulgaris* subsp. *adanensis*, and *B. macrocarpa* are not shown as these species occur along the whole coastline of Western Eurasia (1,10,11,49).

The plants were grown under long day conditions in a greenhouse and were obtained and grown in accordance with German legislation. Genomic DNA was isolated from young leaves using the NucleoSpin ® Plant II protocols from Macherey & Nagel. Each high-quality gDNA (200 ng) was fragmented by sonication using a Bioruptor (Fa. Diagenode) and subsequently used for library preparation with the TruSeq Nano DNA library preparation kit (Fa. Illumina). End repaired fragments were size selected by AmpureXp Beads (Fa. Beckmann-Coulther) to an average size of around 700 bp. After end repair, A-tailing and ligation of barcoded adapters, fragments were enriched by eight cycles of PCR. The final libraries were quantified using PicoGreen (Fa. Quant-iT) on a FLUOstar plate reader (Fa. BMG labtech) and quality checked by HS-Chip on a 2100 Bioanalyzer (Fa. Agilent Technologies). Before sequencing all libraries were pooled depending on the genome size and ploidy of each accession and sequenced 2 x 250 nt on a HiSeq1500 in rapid mode over two lanes using onboard cluster generation. Processing and demultiplexing of raw data was performed by bcl2fastq-v2.19.1 to generate FASTQ files for each accession.

### Plastome assemblies and annotation

Trimmomatic (v0.39) (50) was applied to remove adapter sequences (ILLUMINACLIP:adapters.fa:2:30:10:2:keepBothReads) and to ensure high quality of the reads (SLIDINGWINDOW:4:15 MINLEN:50 TOPHRED33). FastQC (v0.11.9) (51) was used for quality checks. The trimmed reads were subjected to GetOrganelle (v1.7.0) (52) to generate plastome assemblies as suggested for Embryophyta plant plastome sequences (- R 15; -F embplant_pt). The SPAdes (53) kmer settings were set to -k 21, 45, 65, 85, or 105. The contig coverage information and other graph characteristics are used by GetOrganelle to construct the final assembly graphs, which were plotted and visually assessed using Bandage (v0.8.1) (54). The assemblies suggested a circular sequence, however, circular plastome molecules might only comprise a small proportion of all molecules in the cell, whereas other plastome molecules may occur in branched or linear configurations (55–57). The assemblies were submitted in the FASTA format, retaining the possibility to reuse the submitted assemblies as circular or linear sequences. The complex assembly graph of Bmar1 was not automatically resolved. Therefore, single contigs of Bmar1 were sorted manually based on the structure of the other assemblies to enable comparative analyses as described in the following section.

Structural annotation of all plastome assemblies was performed with GeSeq (v2.01) (58). The BLAT (59) search parameters ‘Annotate plastid trans-spliced rps12’ and ‘Ignore genes annotated as locus tags’ were used together with a ‘Protein search identity’ of 25 and a ‘rRNA, tRNA, DNA search identity’ of 85. For HMMER profile search ‘Embryophyta chloroplast (CDS+rRNA)’ was selected and ‘MPI-MP chloroplast references (Embryophyta CDS + rRNA)’ was chosen as reference. The resulting annotation files in the gff format were directly used for further analyses. To avoid confusion, we want to make aware of the fact that *psbN* and *pbf1* are two different names for the same gene.

### Construction of phylogenetic trees

The workflow for the alignment-based phylogenetic analysis is available in Additional file S1G. The position of each gene was extracted from the GFF files obtained through GeSeq. Next, adjacent genes with conserved microsynteny across all investigated samples (including amaranth, quinoa, and spinach as outgroup) were identified and the interleaved intergenic regions of these neighbouring genes were extracted. For overlapping genes, the extraction of an intergenic region was not possible.

Using the gene sequences and intergenic regions of all samples, gene/region specific alignments were performed using MAFFT (v7.299b) (60). High accuracy was ensured using the L-INS-I method. To align sequences with different orientations, the parameter ‘--adjustdirection’ was used. The alignments were trimmed using trimAl (v1.4.rev22) (61). The gap threshold was set to ‘-gt 0.8’, whereas the threshold for the minimum average similarity was set to ‘-st 0.001’. Then, the single alignments were concatenated and the resulting alignment matrix was inspected using SeaView (62). Manual adjustment was not necessary.

RAxML-NG (v1.0.0) (63) was used for ML analysis together with bootstrapping (Model: GTR+FC+G8m). The substitution matrix GTR (for DNA) was applied together with the model parameter ‘G8’ and ‘F’. The parsimony-based randomised stepwise addition algorithm was selected for the starting tree (--tree pars{10}). The number of replicate trees for bootstrapping was set to 200. The resulting tree was visualised using FigTree (v1.4.4) (64).

Location-based clustering of the clades in the tree was performed manually based on the sampling locations (Additional file S2C).

To identify a reduced set of gene sequences and intergenic regions for the construction of a phylogenetic tree distinguishing all accessions, the sequences were iteratively added with increasing alignment identities until all species were separated by informative positions.

To investigate the region dependent phylogeny different additional data matrices were constructed for (I) all gene regions, (II) all intergenic regions, (III) complete coding sequences, (IV) first and second codon position and (V) third codon position only. Therefore, coding sequences for all 79 protein coding genes were extracted from the Genbank annotation files of our plastome assemblies. Start and stop codons were removed and extracted sequences were processed as described above for ML analysis.

To extract the 18s rRNA gene sequence from *B. corolliflora* (Bcor1) as representative of the *Corollinae*, SOAPdenovo2 assemblies were generated using the trimmed reads as input. SOAPdenovo2 (v2.04) (65) was tested with different kmer sizes ranging from 67-127 in steps of 10. The resulting assembly with the highest N50 length was used for further investigations. The reference 18s rRNA gene sequence for *B. vulgaris* subsp. *vulgaris* was retrieved from the NCBI (GeneID=809573). The 18s rRNA gene sequence for Bcor1 was identified via BLAST and then extracted from the SOAPdenovo2 assembly, consecutively adding the following overlapping BLAST hit with the smallest e-value. Next, these extracted sequences were combined for a 18s gene sequence reconstruction. The assembled 18s rRNA gene sequence and the corresponding reference gene sequence were aligned via MAFFT and inspected using SeaView.

### Mitogenome and plastome phylogeny based on kmer-derived phylogenetic splits

SANS serif (v2.1_04B) (66,67), a method based on colored de Bruijn graphs, was selected for the reconstruction of additional phylogenies using variable input data (mitochondrial reads, chloroplastic reads, and full plastome assemblies). This method does not require prior assembly of the reads and is therefore especially suitable for the mitochondrial sequences which could not be fully assembled using GetOrganelle due to the relatively low available sequencing depth and also higher complexity of the mitogenome in comparison to the plastome.

Reads were assigned to the plastome or the mitogenome, respectively, after mapping with BWA-MEM (v0.7.13) (68) against the sugar beet reference genome sequence, including the respective sugar beet chloroplast (KR230391.1) and sugar beet mitochondrial sequence (BA000009.3), which were published independently from this study. This enabled the extraction of reads mapping with higher confidence to the chloroplast/mitochondrial sequence in contrast to mapping to the nucleome (with e.g. a few mismatches) and *vice versa*. Therefore, ‘samtools view’, with the -b and -h options, was used after indexing the BAM file. The resulting BAM file was then further processed using ‘samtools collate’ and converted to the FASTQ format to extract the corresponding reads (chloroplast or mitochondria, respectively) using ‘samtools fastq’. After that, for the read-derived phylogenetic analyses, a colored de Bruijn graph was constructed using Bifrost (v1.0.5) (69) to filter kmers which only occur once in the dataset. This graph was then used as input for SANS serif. In addition, a phylogeny was reconstructed using the newly constructed plastome assemblies as direct input for SANS serif.

Different parameters were applied to test the robustness of the results. These arguments include different mean weight functions (-m; geom vs. geom2), the number of splits in the output list (-t; all vs. 10n) as well as various kmer sizes (11, 21, 31 and 61). The SANS serif output file was then converted to the nexus format (sans2nexus.py) and subsequently visualised using Splitstree5 (70). To analyse the discrepancies between trees derived from different methods, SANS serif was used with the option ‘strict’ to generate an output file in the newick format which was then visualised using FigTree (64). Further, the SANS serif script ‘comp.py’ was used to calculate weighted (length of the edges/size of the splits is taken into account) precision and recall (combined in F1 scores) while using each tree as reference/ground truth in an ‘all vs. all’ comparison. In this use case, precision means ‘the total weight of all correctly predicted splits divided by the total weight of all predicted splits’. Further, weighted symmetric set distances and Robinsons-Foulds distances were calculated for each comparison. Details can be found in Additional file S1D. For this analysis, the trees constructed with different input data (cp_reads, mt_reads, and cp_assemblies) and the fixed parameters ‘-m geom2, -t 10n, -k 31’ were compared.

### Investigation of alignment identities

The alignment identities for each plastome gene sequence or intergenic region were calculated to infer the phylogenetic information of all sequences. The events (SNV or InDel) were detected by iteration over each position in the sequence. The identity score (percent identity) was calculated by division of conserved positions (number of residues [position in alignment] with the same nucleotide in all accessions) by the number of residues in the alignment (’sequence length’). Alignment identities were calculated (i) for all accessions and (ii) for all accessions including outgroup reference plastome sequences (amaranth, quinoa, and spinach). Visualisation of the results was performed using matplotlib (71). Next, potential hotspots for SNVs and InDels in the plastome sequences were investigated.

## Declarations

### Ethics approval and consent to participate

The material of the KWS Saat SE, Einbeck, and IPK Gatersleben was transferred under the regulations of the standard material transfer agreement (SMTA) of the International Treaty. Plants were grown in accordance with German legislation

### Consent for publication

Not applicable.

### Availability of data and materials

Sequence reads have been submitted to the European Nucleotide Archive (ENA; Additional file S2D). The plastome assemblies and the corresponding annotations are available at ENA/GenBank (PRJEB45680). Additional information used in phylogenetic analyses are included in the supplementary files.

### Competing interests

The authors declare that the research was conducted in the absence of any commercial or financial relationships that could be construed as a potential conflict of interest.

### Funding

Open Access funding enabled and organized by Projekt DEAL. KS is funded by Bielefeld University through the Graduate School DILS (Digital Infrastructure for the Life Sciences).

### Authors’ contributions

BP, BW, TH and DH designed the study. NS selected and cultivated the plants and performed DNA extraction. PV designed the layout for sequencing. BP and KS developed and implemented the bioinformatic methodology. KS analysed the data and prepared the figures and tables. KS, BP, NS and TH wrote the manuscript. All authors read and approved the final manuscript.

## Acknowledgements

We acknowledge support for the Article Processing Charge by the Deutsche Forschungsgemeinschaft (German Research Foundation) and the Open Access Publication Fund of Bielefeld University. We thank the CeBiTec Bioinformatic Resource Facility team for great technical support and Dr. Roland Wittler for great support with the SANS serif software.

## Supplementary Material

Additional file S1A: Geographic distribution of the Betoideae species as described in the literature.

Additional file S1B: Distribution of coverage values and assembly length (in bp) for each region and the total assemblies.

Additional file S1C: Circular and linear plots of selected plastome assembly sequences. Additional file S1D: Distance metrics for the comparison of splitstree results.

Additional file S1E: Reduced phylogenetic tree based on 53 gene and intergenic regions. Additional file S1F: Phylogenetic trees based on different sequence matrices.

Additional file S1G: Workflow for the construction of phylogenetic trees.

Additional file S2A: Sequence identities [%] of all investigated plastome gene sequences and intergenic regions including outgroup reference sequences.

Additional file S2B: Sequence identities [%] of all investigated plastome gene sequences and intergenic regions excluding outgroup reference sequences.

Additional file S2C: Accession IDs, taxonomy, read and assembly statistics and geographic location of the investigated accessions of the Betoideae subfamily.

Additional file S2D: SRA-IDs of the processed read datasets.

## Notes

### Competing Interest Statement

The authors have declared no competing interest.

### Summary of Updates

minor changes - added titels to all additional files

