## Additional file S1 for "Complete pan-plastome sequences enable high resolution phylogenetic classification of sugar beet and closely related crop wild relatives"

### S1A: Geographic distribution of the Betoideae species as described in the literature.

| species | geographic distribution | references |
|---|---|---|
| *B. macrocarpa* | distributed along the coastal regions of Portugal, Spain, Greece, Italy, North-West Africa (Morocco and Algeria), Turkey and Israel, but can also be found on the Canary Islands; tetraploid *B. macrocarpa* is only found in Portugal and the Canary Islands | (Buttler, 1977; Kadereit et al., 2006; Frese et al., 2011; Romeiras et al., 2016; Touzet et al., 2018) |
| *B. vulgaris subsp. adanensis* | distributed in the eastern Mediterranean area, including Greece, Cyprus, Turkey and Syria | (Romeiras et al., 2016; Touzet et al., 2018) |
| *B. patula* | endemic to Madeira Island and the Desertas Islands | (Frese et al., 2011; Romeiras et al., 2016; Touzet et al., 2018) (Figure 1) |
| *B. vulgaris subsp. vulgaris* | widely cultivated and just like *B. vulgaris subsp. maritima* spread along the Atlantic coast of Western Europe and the Mediterranean coast, including North Africa, Macaronesia and Western Asia | (Romeiras et al., 2016; Touzet et al., 2018) |
| *B. vulgaris subsp. maritima* | spread along the Atlantic coast of Western Europe and the Mediterranean coast, including North Africa, Macaronesia and Western Asia | (Romeiras et al., 2016; Touzet et al., 2018) |
| *B. corolliflora* | distributed in South-West Asia and the Caucasus | (Kadereit et al., 2006; Romeiras et al., 2016) |
| *B. intermedia* | distributed in Armenia and Turkey | (Kadereit et al., 2006; Romeiras et al., 2016) |
| *B. lomatogona* | distributed in Turkey, Iran, Azerbaijan and Armenia | (Kadereit et al., 2006; Romeiras et al., 2016) |
| *B. macrorhiza* | distributed in Armenia, Azerbaijan, Dagestan and Turkey which roughly corresponds to the area of the Caucasus as well | (Kadereit et al., 2006; Romeiras et al., 2016) |
| *B. nana* | endemic to the Greek mountains | (Kadereit et al., 2006; Frese et al., 2011; Romeiras et al., 2016) (Figure 1) |
| *P. patellaris* | distributed on the Canary Islands, the Iberian peninsula, the Balearic Islands, in Italy, North Africa (Algeria and Morocco), Madeira and Cape Verde | (Kadereit et al., 2006; Frese et al., 2011; Romeiras et al., 2016) (Figure 1) |
| *P. procumbens* | restricted to the Canary Islands, Madeira, and Cape Verde | (Kadereit et al., 2006; Frese et al., 2011; Romeiras et al., 2016) (Figure 1) |
| *P. webbiana* | even more restricted as this species can only be found on Gran Canaria (and possibly Tenerife and Fuerteventura) | (Kadereit et al., 2006; Frese et al., 2011; Romeiras et al., 2016) (Figure 1) |

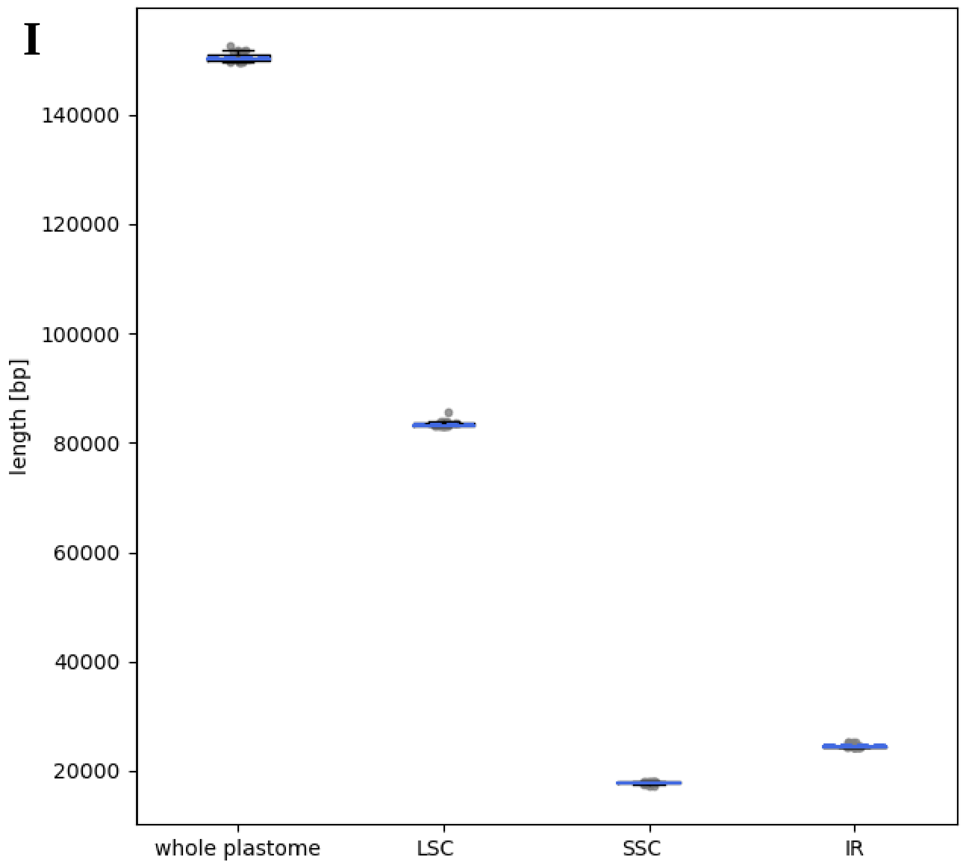


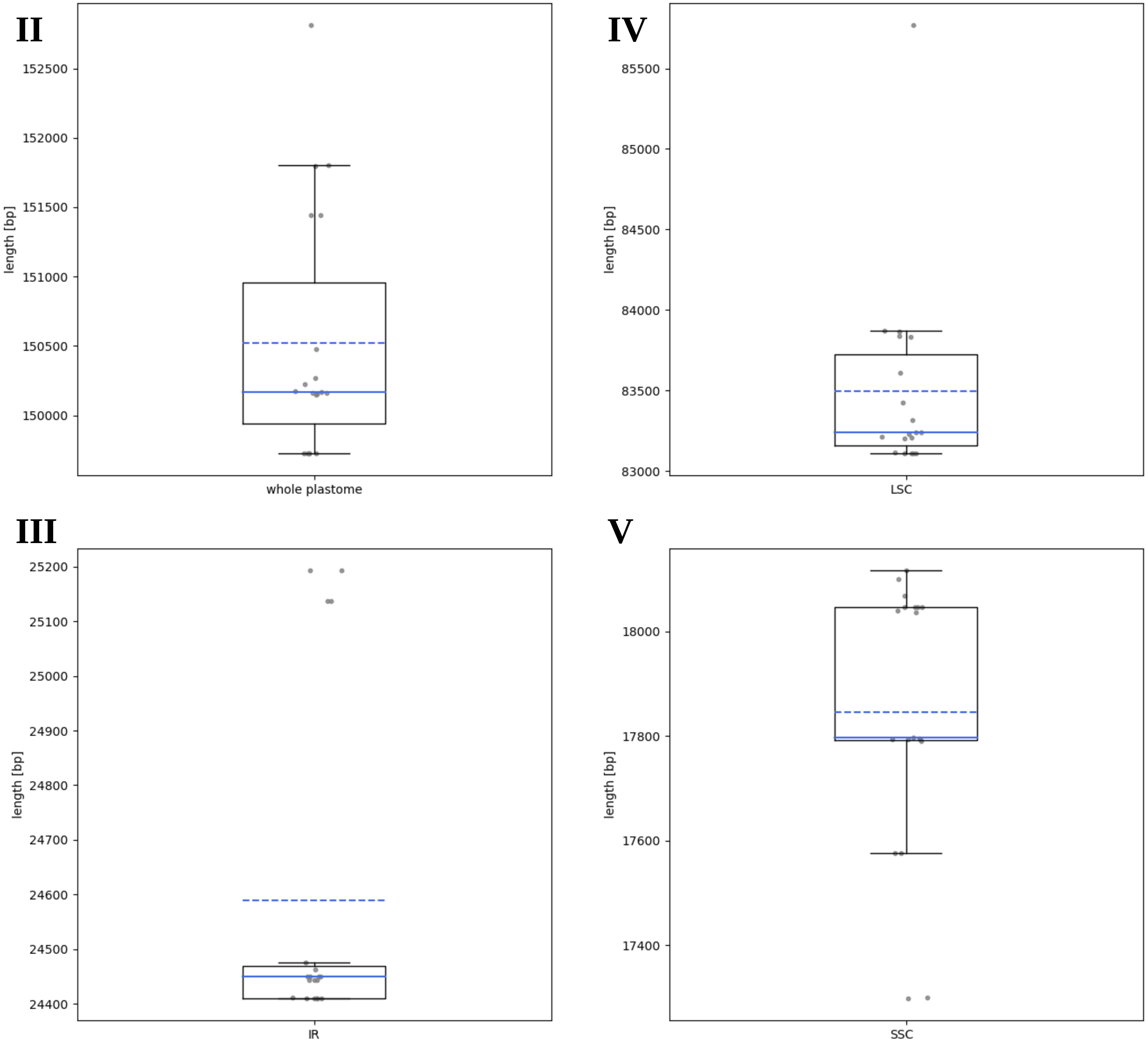


**S1B: Distribution of the length [bp] of different regions of the plastome assemblies.** I) Overview of all plastome regions. II-V) Detailed view of the length of the whole plastome, IR, LSC and SSC, respectively.

VI

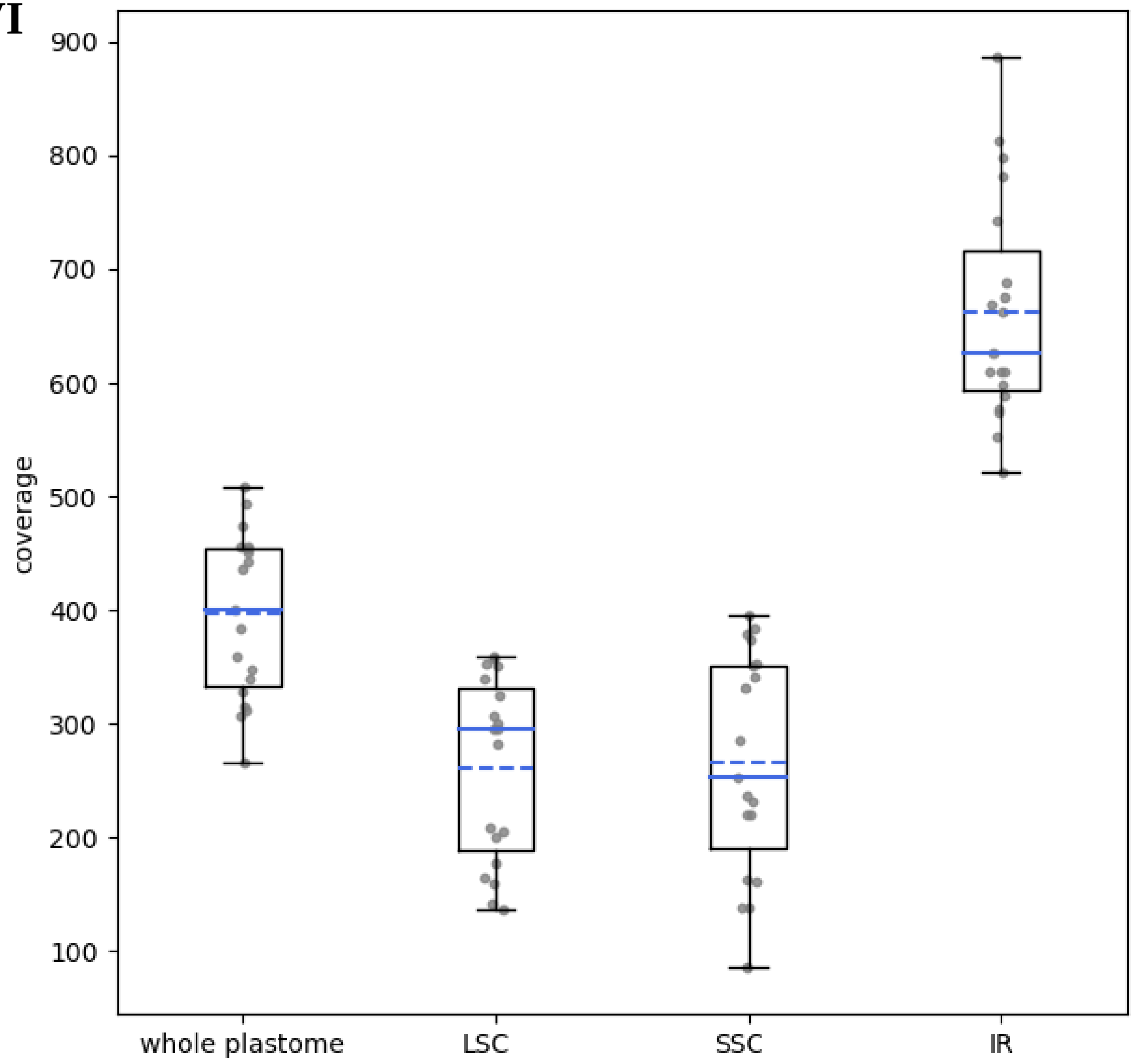


**S1B - VI: Distribution of the coverage of different regions (whole plastome, LSC, SSC and IR) of the plastome assemblies.**

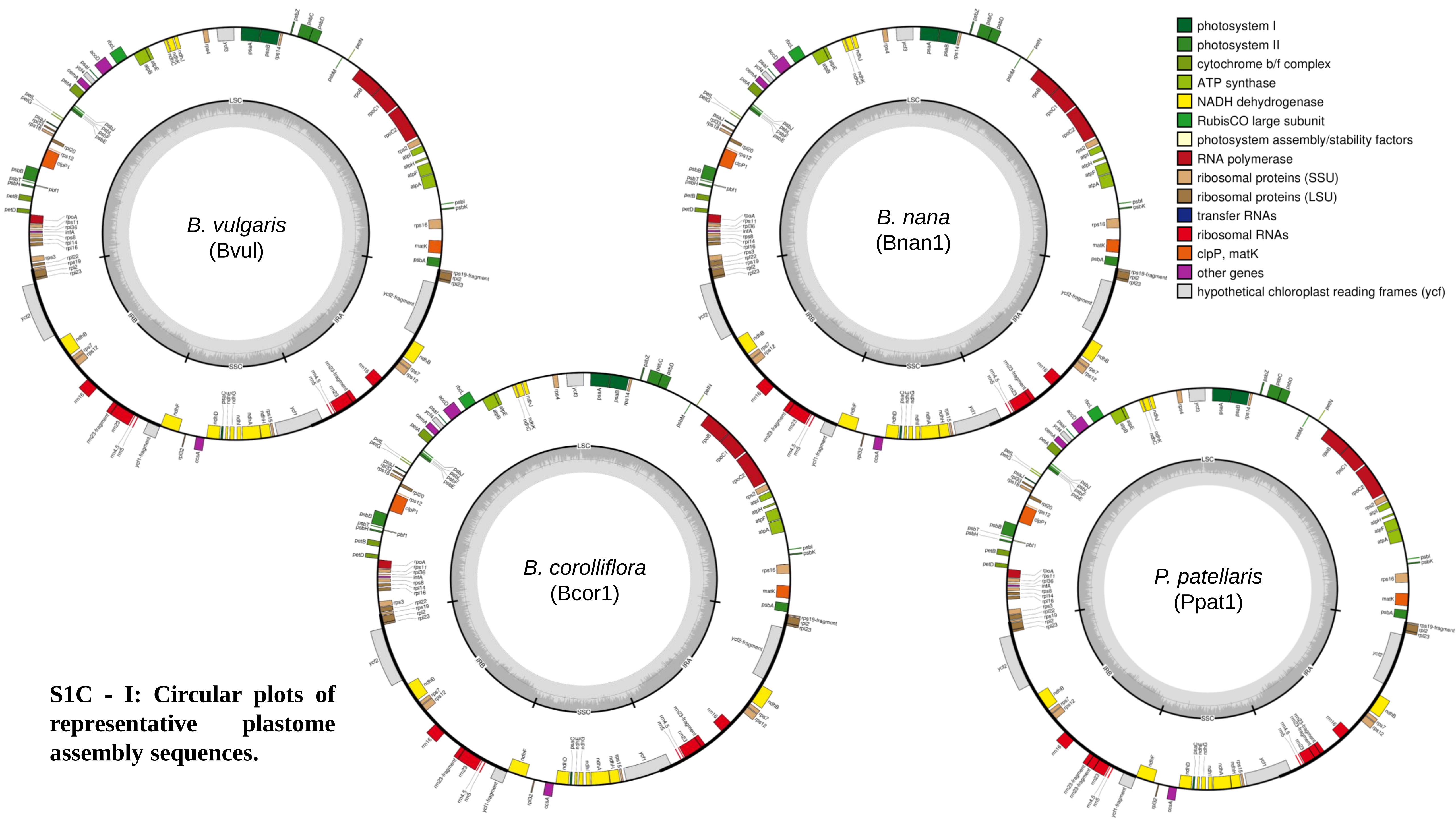


**S1C - I: Circular plots of representative plastome assembly sequences.**

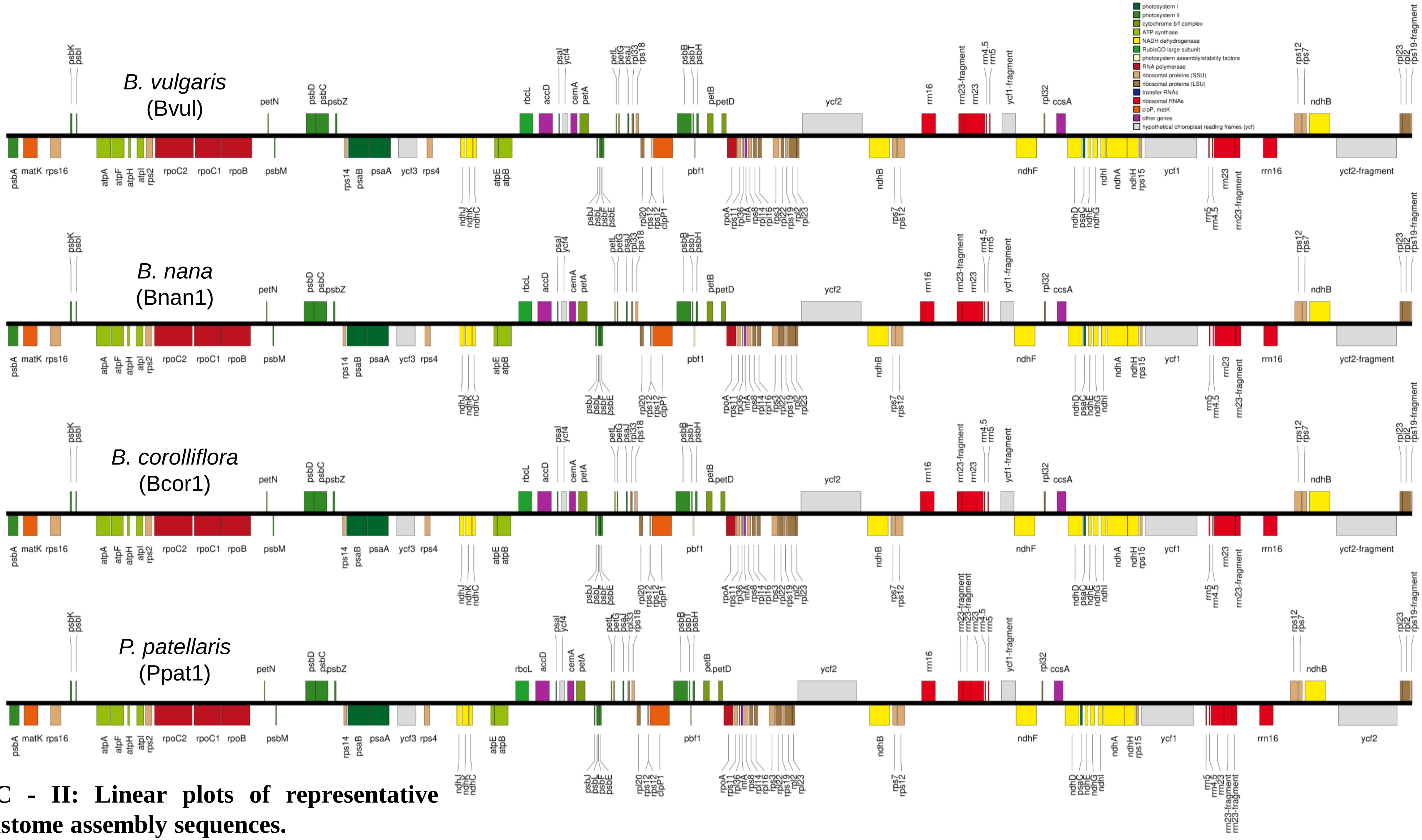


**S1C - II: Linear plots of representative plastome assembly sequences.**

**S1D: Distance metrics for the comparison of splitstree results.**

| Comparison | F1 score | Weighted symmetric set distance | Robinsons-Foulds distance |
|---|---|---|---|
| cp_reads vs. mt_reads | 0.94 | 0.110 | 48 |
| cp_reads vs. cp_assemblies | 0.90 | 0.177 | 52 |
| mt_reads vs. cp_assemblies | 0.91 | 0.163 | 62 |

$$\text{F1 score} = 2 * \frac{\text{precision} \ * \text{recall}}{\text{precision} + \text{recall}}$$

$$\text{Weighted symmetric set distance} = \frac{\text{total weight of splits}(\text{A, not in B})}{\text{total weight}(\text{A})} + \frac{\text{total weight of splits}(\text{B, not in A})}{\text{total weight}(\text{B})}$$

$$\text{Robinsons}-\text{Foulds distance} = \text{sum of splits in A, not in B} + \text{sum of splits in B, not in A}$$

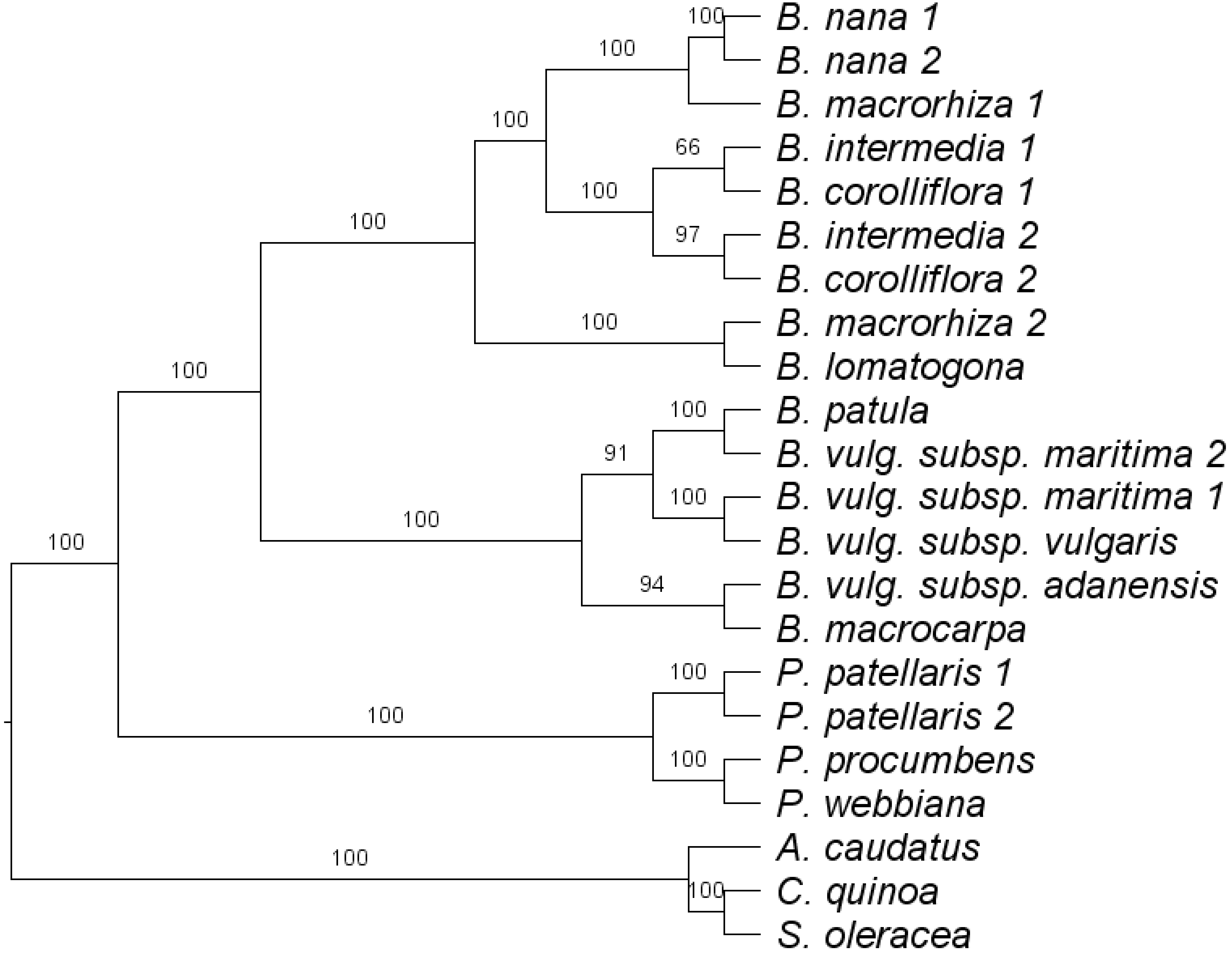


**S1E: Phylogenetic tree based on the diagnostic set of 53 gene sequences.** The plastome sequences of three Caryophyllales species (amaranth, quinoa and spinach) were used as outgroup. Bootstrap support values are shown above each branch. The resulting phylogeny is based on the variation in 53 sequences from the plastid genome of 18 accessions and species (plus outgroup).

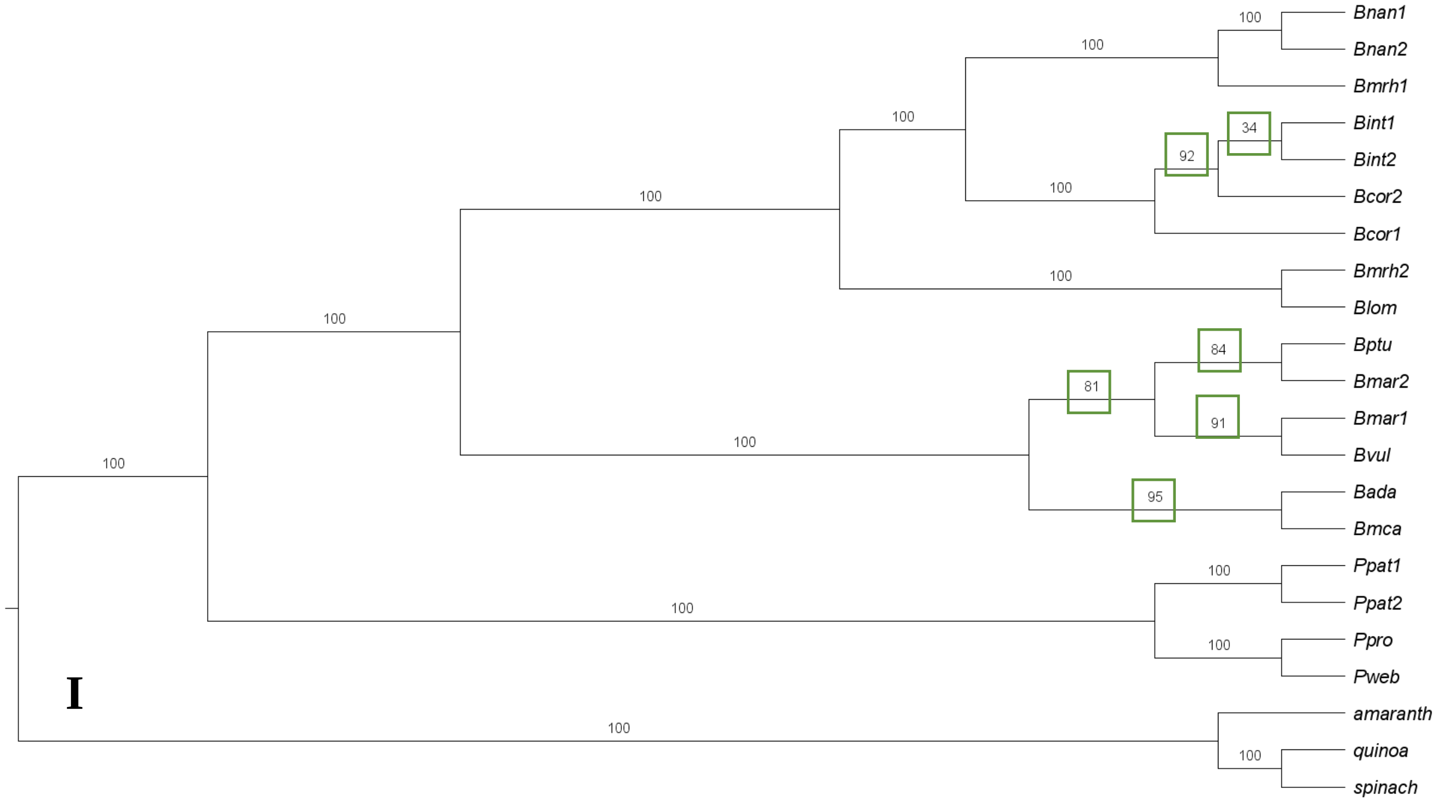


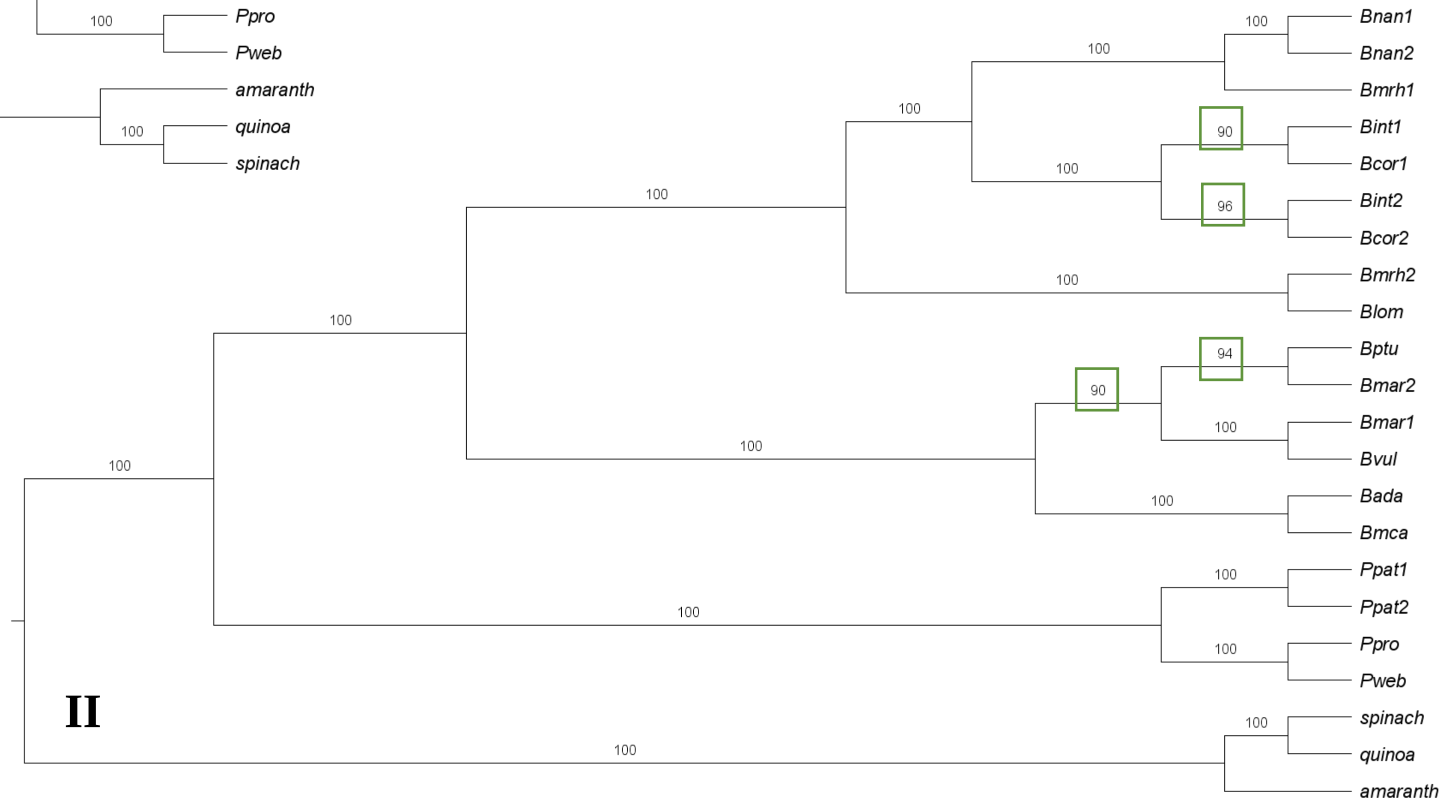


**S1F: Phylogenetic trees based on different sequence matrices.** I) gene regions; II) intergenic regions. Bootstrap values are shown above each branch. Trees are derived from ML analysis with RAxML-NG with 200 bootstrap replicates. Bootstrap values <100 are marked in green.

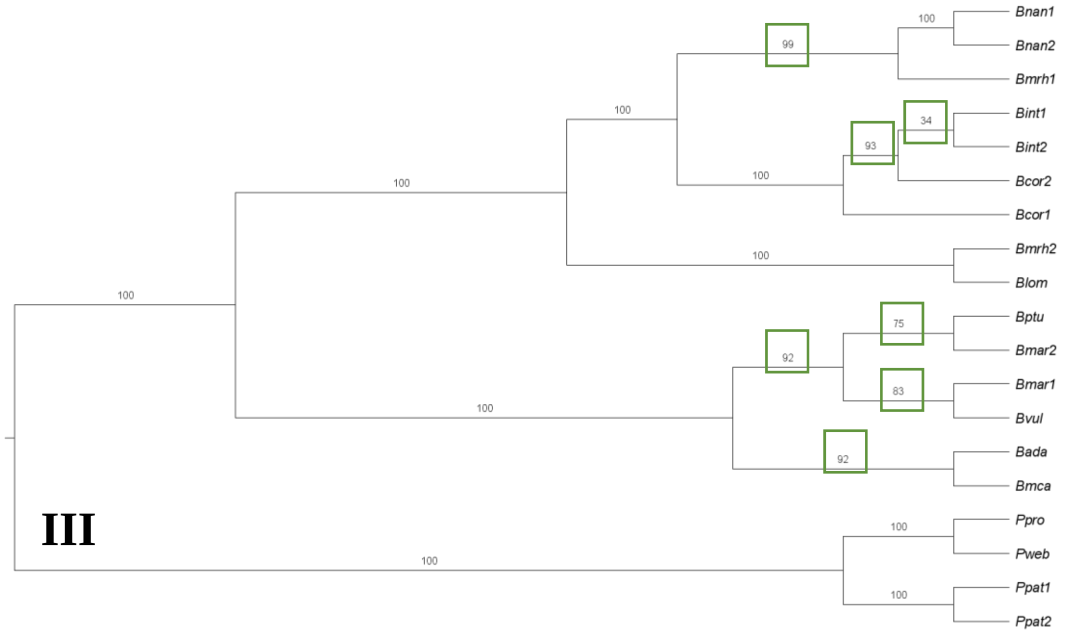


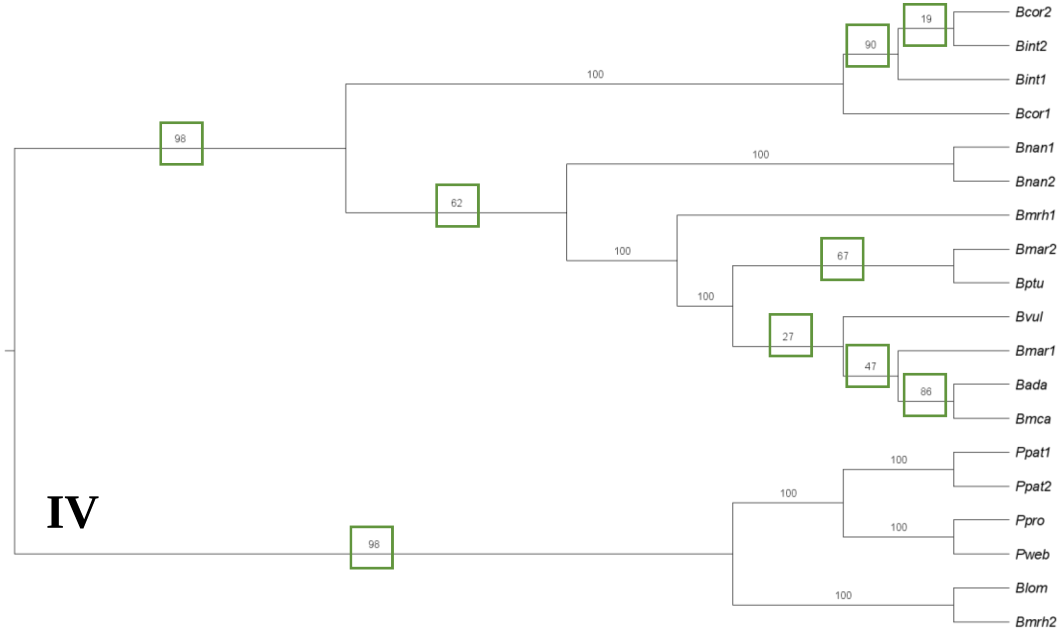


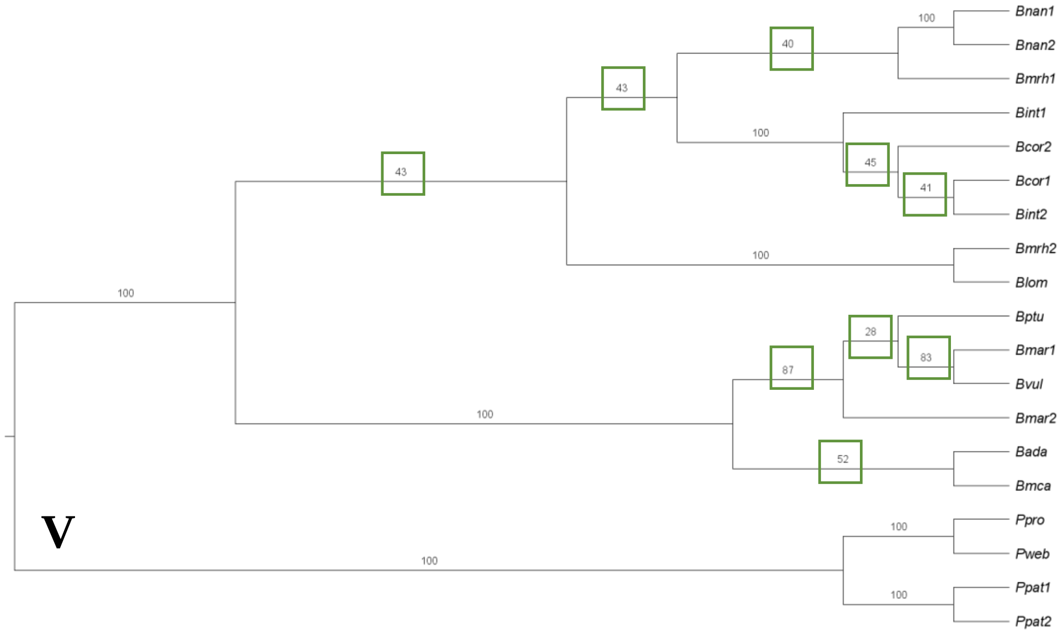


**S1F: Phylogenetic trees based on different sequence matrices.** III) coding sequences; IV) first and second codon positions; V) third codon position. Bootstrap values are shown above each branch. Trees are derived from ML analysis with RAxML-NG with 200 bootstrap replicates. Bootstrap values <100 are marked in green.

Annotation (GFF files) + Assembly (FASTA files)

Extract position of each gene per accession from annotation file

Identify genes which are located next to each other in each accession (including outgroup plastome sequences)

Extract intergenic regions of these neighboring genes from annotation and assembly files

Extract gene sequences from annotation and assembly files

Gene/region specific alignments using MAFFT (v7.299b)

Trimming of each alignment using trimAI (v1.4.rev22)

Concatenation of all single alignments

RaxML-NG (v1.0.0) for ML and bootstrapping analysis

Visualization of the phylogenetic tree using FigTree (v1.4.4)

**S1G: Workflow for the construction of phylogenetic trees.**
